# Parsimonious Creatures: Building Gene Regulatory Networks Robust to Somatic Evolution

**DOI:** 10.64898/2026.11.07.764957

**Authors:** Hyrum Diesen, Frederick R. Adler

## Abstract

For early multicellular life to have formed and maintained cellular organization, preventing loss of controlled cell replication due to somatic evolution was of critical importance, and remains important to this day for control of cancer. We study how gene regulatory networks (GRNs) can be designed to slow the divergence of somatic clonal lineages in the context of early multicellular life. We model the mutation of GRNs within simplified 12-celled organisms we term “parsimonious creatures”, who must replicate from a single cell to 12 cells and then prevent replication past 12 cells to maximize fitness. We found two minimal motifs can achieve these goals, but both are prone to breakdown through somatic evolution: The feedforward motif, which is loosely regulating but relatively robust, and the simple step motif, which uses strong interactions and feedback to form memoryless ultrasensitivity, but is fragile. These simple motifs may be duplicated to form redundant networks via one of two structures: averaging redundancy, which averages multiple sources of information, and min redundancy, which adopts the most conservative source. Averaging redundancy uniformly slows somatic evolution, while min redundancy stalls somatic evolution until neutral mutations accumulate. Either may be superior dependent on life history. However, averaging redundancy slowly increases fitness as the network becomes larger, but large min redundancy networks eventually become “paralyzed”, increasing the probability of senescence and reducing fitness. These GRN designs are idealized, yet provide imperfect solutions with fundamental tradeoffs, offering insight as to why somatic evolution remains an incompletely solved problem for multicellular life.

## Introduction

Multicellular life relies on coordination of cell replication rates among cells to maintain proper organ and body size. Successful coordination depends on signals between cells being correctly interpreted and acted upon by individual cells [1]. However, if individual cells mutate to divide more quickly, the inexorable logic of natural selection will lead them to increase in frequency. This somatic evolution can lead to runaway hyperplasia to the detriment of the organism’s fitness [2, 3]. For multicellular life to persist, the conflicting levels of selection within and among organisms must be resolved [4]. In organisms with a segregated germline, somatic cells die when the organism dies, favoring the germline to encode mechanisms to maintain proper tissue size.

Nonetheless, in the case of cancer, a class of diseases under the broader umbrella of somatic evolution, we frequently observe cancer as a cause of death in individuals, even at around 5 percent in species such as elephants thought to be highly resistant to cancers at the population level [5]. Measuring population level fitness loss from spontaneous cancers in natural environments remains difficult. Nevertheless, current studies show that cancer prevalence (rather than fitness loss) to be supported by a model of stabilizing selection, and is highly variable depending on the species [6]. Moreover, hyperplasia is particularly prevalent in some populations, present post-mortem in a staggering 35 and 65 percent of wild opossums and ferrets respectively. We can conclude that mechanisms have been honed by selection to reduce, but not eliminate, cancer.

Several solutions to this problem have been observed in multicellular life. First, and most simply, organisms can reduce the somatic mutation rate, such as through DNA repair [7]. This delays, but does not remove the challenge; once mutations have occurred, other mechanisms are needed to prevent invasion by hyperplastic mutants. Second, intercellular controls can detect, inhibit, and eliminate cells that divide inappropriately. These mechanisms are widespread in long-lived metazoans, and require mechanisms for communication and coordination which are themselves subject to mutational breakdown [8]. We here focus on a third mechanism, intracellular robustness through design of the systems that control cell replication.

Cell replication is controlled by gene regulatory networks (GRNs). The effects of mutations are filtered through these networks. For example, a gene regulatory network could be robust to somatic evolution if it minimized the effects of mutations on cell replication rates, biased somatic mutants toward lower replication, or if it somehow detected a mutation and then toggled outputs to select against its own lineage. Engineers can build complex devices with only a few backup systems, such as triple redundancy in aerospace computers or failsafe mechanisms in escalators and elevators, that reduce the chance of catastrophic failure to the point it is never observed on practical timescales. In contrast, multicellular life has existed on Earth for at least 600 million years [9], with a concommitant increase in gene count and overall genome size [10], and yet multicellular life still does not display this level of robustness to potentially fatal somatic evolution.

There are three broad reasons for this apparent failure. First, these mechanisms might be costly in terms of energy use and growth rate, and therefore likely to reduce reproduction in young organisms. Second, a highly effective system will have redundant pathways to deal with rare cases, and therefore suffer from the “paradox of robustness” because selection to maintain redundant pathways is weak [11]. Third, in the words of Michael Crichton, “life finds a way”; there may simply be constraints on the potential of any GRN design to be robust to somatic evolution due to natural selection’s ability to find its way over almost any barrier.

We address this last possibility by attempting to build a counterexample. We develop a set of theoretical models designed to discover GRNs that regulate cell replication and are highly robust to somatic evolution. To this end, we model the lives of “parsimonious creatures”: minimal in-silico organisms with life histories intended to mimic early multicellular lifeforms with gene regulatory networks experiencing somatic evolution. These organisms begin life from a single cell, and grow to reproduce when the number of cells is close to an optimal number *N*^*\**^ in the face of cell death, organism death, and somatic evolution that can drive the number of cells past *N*^*\**^.

We have combined a human design approach with a variety of established and custom optimization methods to discover GRN designs that are highly robust to somatic evolution in parsimonious creatures. We present three key results.

1. We show two minimal motifs that achieve and maintain the optimal size in the absence of somatic mutation, but differ in tightness of regulation and robustness to mutation.
2. We use these motifs as building blocks to create two forms of redundancy that are better at maintaining the optimal number of cells in the face of somatic mutation.
3. We show fundamental tradeoffs of each form of redundancy, and how these tradeoffs lead to one form being favored over the other depending on the life history parameters and network size.

## Results

For our primary results, we assume that the soma is fully segregated from the germline, and we will ignore the germline entirely. parsimonious creatures will therefore be tested completely independently of each other with a gene regulatory network that is predefined at the beginning of it’s life rather than one that is inherited from another organism.

To regulate cell number, each cell should be regulating it’s own rate of replication, *λ*, in response to the number of cells currently in the organism, *N*. In the gene regulatory network of each cell, *N* will act as a signal that will interact with transcription factors at rate 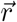 by increasing their degradation rate. These transcription factors will inhibit each other in a potentially complex network **K** by competing for promoter sites, but will also may inhibit the cell’s replication rate *λ* at rate 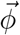. We will assume that all interactions in a GRN are inhibitory and all nodes (TFs and *λ*) are fully expressed by default. This GRN will form the cell’s replication function, *λ*(*N*). We assume that cells cannot control their own death rates, and cell death rate *δ*_*C*_ is constant.

Therefore, to maintain a cell number close to the reproductive optimum *N*^*\**^ (which we initially set to be *N*^*\**^ = 12) the replication function needs to be decreasing: large when *N < N*^*\**^ and small when *N > N*^*\**^. To quantify how well *N* is constrained, we created a deterministic model of parsimonious creatures: life begins with a single somatic cell with a some GRN that produces some *λ*(*N*). Using master equations, organisms can be at any *N*, and can transition to higher or lower *N* with probabilities defined by *λ*(*N*), while organism death transitions an organism to be “reborn” back to *N* = 1, and we set absorbing boundary conditions at *N* = 50. Exact equilibrium probabilities of each *N* can thus be solved for equilibrium to quantify how probable it is for the number of cells in the organism to be close to *N*^*\**^. This deterministic model of parsimonious creatures enabled us to rapidly and exactly test how well a GRN will keep *N* constrained around *N*^*\**^.

However, this deterministic model must assume the absence of somatic mutation, which will not allow us to test whether a GRN is robust to somatic evolution in parsimonious creatures. Therefore, we created an additional stochastic simulation-based model of parsimonious creatures. parsimonious creatures begin life with a single somatic cell with a predefined GRN. This single cell will stochastically perform a division event at rate *λ*(1), and daughter cells will inherit this GRN, which will then solve to *λ*(2), and so on. To include somatic mutations, any mutateable interaction in the GRN of any cell may mutate at rate *µ* to become stronger or weaker at mutation magnitude *σ*. Cells may therefore inherit a GRN which forms a perturbed *λ*(*N*). Ultimately, a more aggressive *λ*(*N*) will naturally cause a lineage to be favored in the organism over time, and will eventually drive *N* to become larger.

### The feed-forward motif

The smallest possible gene regulatory network that has at least one transcription factor and generates a non-constant replication function *λ*(*N*) has a single TF which inhibits replication, but is also inhibited by *N* via degradation. However, we have constrained our GRNs to purely inhibitory interactions, and such a network produces an increasing replication function of *N*, and this would not keep *N* constrained. Therefore, at least two TFs are necessary to produce a decreasing replication function by “inhibiting an inhibitor”: a signal-receiving TF that inhibits a downstream TF which in turn inhibits replication. We refer to this minimal motif which keeps *N* constrained as a “feedforward motif”.

We then optimized the weights of interactions in the feedforward motif in our deterministic model of parsimonious creatures. Blackbox optimization [12] found weights where *N* is optimally constrained around *N*^*\**^. Regardless of the strength of interactions in this topology, the shape of the resulting replication function is gradual (Fig. 1). Therefore, the cell number in an organism over time would have a wide distribution around *N*^*\**^ which results in relatively low fitness. However, when we consider somatic mutations in our stochastic simulation based model of parsimonious creatures, we find that this simplicity grants the feed forward motif robustness against somatic mutation (Fig. 1).

**Figure 1:**
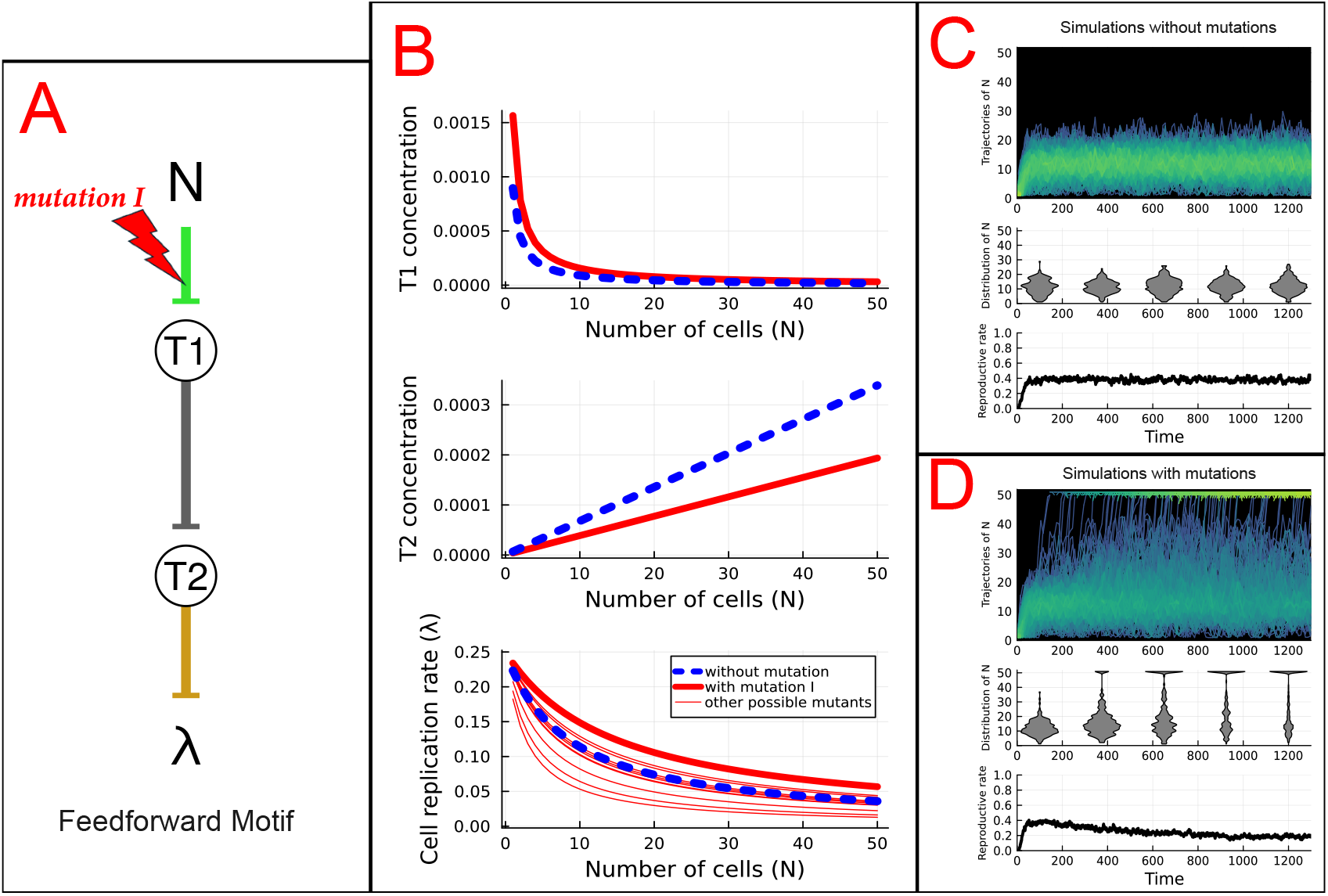
The feedforward motif’s topology, it’s resulting cell replication function and it’s behavior in simulations of parsimonious creatures with and without somatic mutation. (A) The topology of feedforward motif is shown, where *N*, the number of cells in the organism, acts as a signal acting on a GRN formed from the transcription factors *T*_1_ and *T*_2_ to regulate the cell’s replication rate, *λ*. An example mutation *I* is illustrated with a red lighting symbol modifying the rate at which *N* inhibits *T*_1_. (B) *T*_1_, *T*_2_, and *λ* are plotted as a function of *N* prior to mutation in blue. *T*_1_, *T*_2_, and *λ* with respect to *N* are also plotted in red where the example mutation *I* illustrated in A has been applied. *λ*(*N*) is also plotted in red for other possible mutations sampled in the motif. (C, top) 200 parsimonious creatures were independently simulated with the feedforward motif in the absence of both mutation and organism death, where trajectories of *N* are plotted over time in each parsimonious creature. Overlapping trajectories are plotted brighter to illustrate density. (C, middle) The distribution of *N* among all simulations of parsimonious creatures over time are shown as violin plots. (C, bottom) The average reproductive rate of all simulations of parsimonious creatures is plotted over time. (D) Simulations of parsimonious creatures with data collected as in C, but where somatic mutations have been introduced.

### The simple step motif

To maximize lifetime reproduction for parsimonious creatures, each cell should grow as fast as possible when the organism is below *N*^*\**^, then stop all growth when the organism is *N*^*\**^ or more cells. Mathematically, each cell should follow a replication function that is a step function with threshold *η* that is equal to *N*^*\**^. To find the smallest motif that produces a step-like replication function, we performed blackbox optimization with the objective as expected fitness from a deterministic model of parsimonious creatures in the absence of somatic mutation. We constrained network size to have fewer than four transcription factors to reduce the size of our search. This revealed a motif with only two transcription factors to be the simplest network to produce a sharp step function (Fig. 2). This simple two TF system has only one feedback and is algebraically solvable for equilibrium as a quadratic (supplemental).

**Figure 2:**
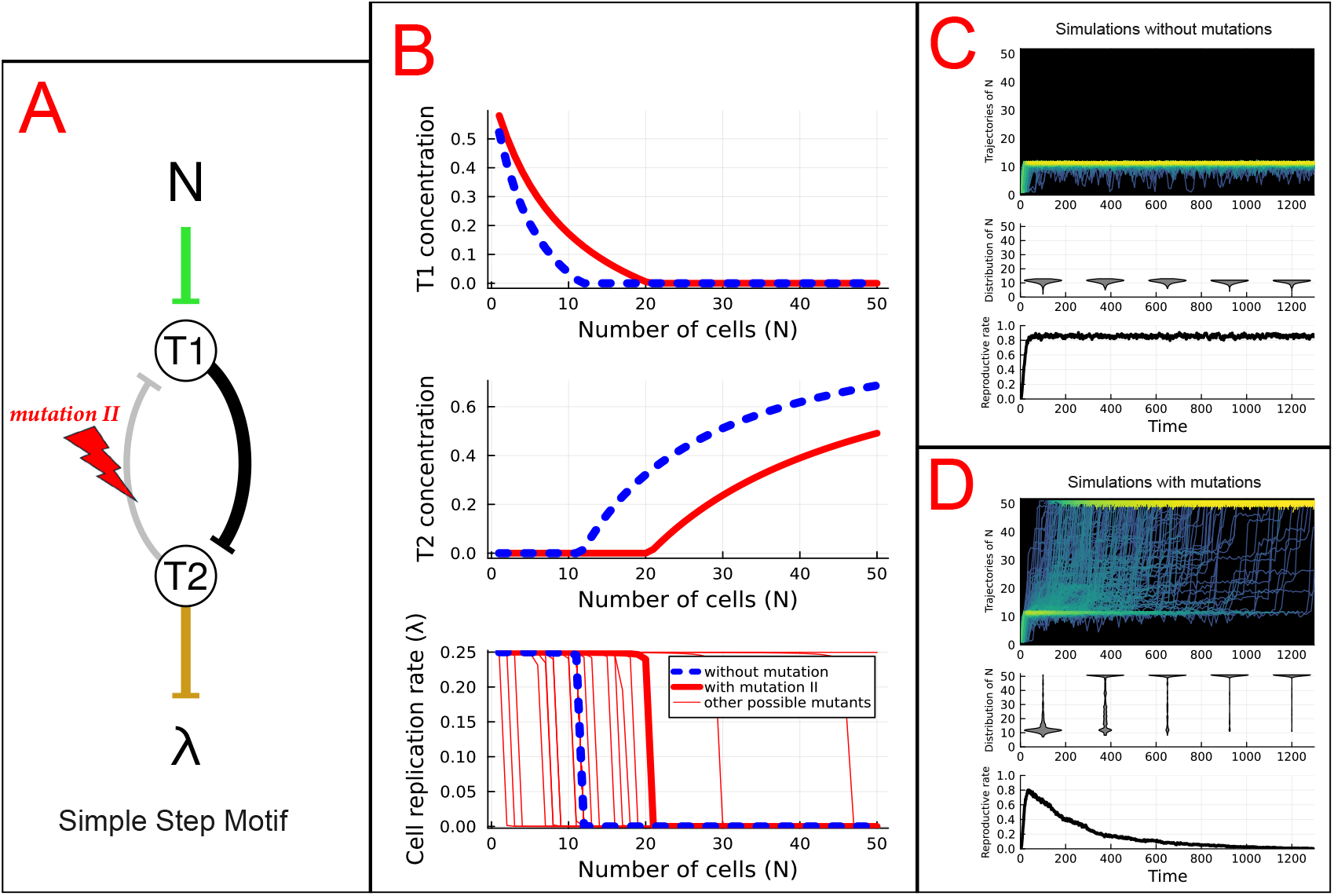
The simple step motif’s topology, it’s resulting cell replication function and it’s behavior in simulations of parsimonious creatures with and without somatic mutation. (A) The topology of the simple step motif is shown, where *N*, the number of cells in the organism, acts as a signal acting on a GRN formed from the transcription factors *T*_1_ and *T*_2_ to regulate the cell’s replication rate, (*λ*). An example mutation *II* is illustrated with a red lighting symbol modifying the rate at which *T*_2_ inhibits *T*_1_. (B) Expression of *T*_1_, *T*_2_, and *λ* are plotted as a function of *N* prior to mutation in blue. Expression of *T*_1_, *T*_2_, and *λ* with respect to *N* are also plotted in red where the example mutation *II* illustrated in A has been applied. *λ*(*N*) is also plotted in red for other possible mutations sampled in the motif. (C, top) 200 parsimonious creatures were independently simulated with the simple step motif in the absence of both mutation and organism death, where trajectories of *N* are plotted over time in each parsimonious creature. Overlapping trajectories are plotted brighter to illustrate density. (C, middle) The distribution of *N* among all simulations of parsimonious creatures over time are shown as violin plots. (C, bottom) The average reproductive rate of all simulations of parsimonious creatures is plotted over time. (D)Simulations of parsimonious creatures with data collected as in C, but where somatic mutations have been introduced.

In the absence of mutation, the simple step motif (SSM) with threshold *η* equal to *N*^*\**^ achieves near maximal lifetime fitness for parsimonious creatures. The sharpness of this threshold increases, as does the fitness, when the maximum value of interaction strength *K* approaches infinity. For our analysis, we set a high upper bound to allow us to create strong step functions with the simple step motif: *K*_*max*_ = 10^8^.

By naïvely optimizing all parameters independently across small gene regulatory network topologies to maximize the fitness in the deterministic model of parsimonious creatures, we discovered a motif which forms a step function, a property commonly referred to as ultrasensitivity. This network does not have hysteretic properties, as the equilibrium states are algebraically solvable to only a single nonnegative solution. This motif achieves these two properties utilizing purely Michaelis-Menten interactions and a signaling term that increases the degradation rate of a single transcription factor. The tradeoff is that this ultrasensitivity requires strong molecular interactions with large values of *K*. While many biological mechanisms and/or motifs displaying ultrasensitivity have been modeled in the literature [13–16], we believe this particular mechanism has not yet been described.

### Building GRNs with redundant motifs to resist somatic evolution

The feedforward motif has relatively low fitness when mutation rate is zero, but is robust to somatic evolution. The simple step motif with a sharp threshold at *η ≈ N*^*\**^ maximizes fitness when mutation rate is zero, but is comparatively fragile and rapidly loses fitness to somatic evolution as mutation rate increases (Fig. 2). We next show that we can utilize multiple redundant motifs to decrease this fragility in multiple ways.

### Averaging redundancy

Gene duplications have been shown to increase mutational robustness of gene regulatory networks by adding redundancy [17, 18] and that redundancy is a fundamental component of robustness [19]. Furthermore, evolutionary genomic evidence often shows long term preservation of redundant duplicates rather than subfunctionalization [20, 21]. Nevertheless, evidence points against gene duplications driving the wholesale formation of motifs in gene regulatory network evolution [22, 23]. However, the simple two-TF motifs we describe could plausibly duplicate jointly as adjacent sequences, or via whole genome duplication, the latter of which has been correlated with motif evolution and is associated with GRN robustness [24–26].

A complete duplication of all the transcription factor genes in a motif would produce a dense matrix of mutual connections between homologous TF genes [27] (Fig. 3, Fig. S6), which may alter function. In the case of the simple step motif, we found the threshold of the step-like replication function remains the same after a complete duplication of the motif. Given that affinities will stay constant immediately after a duplication, the threshold of the step function is nearly identical, but it will become sharper due to increased mutual inhibition, tightening the distribution of *N* to keep it more effectively constrained. For mathematical convenience, we have opted to normalize for this increased inhibition. This correction maintains an identical nonmutant replication function regardless of number of times the motif is duplicated.

**Figure 3:**
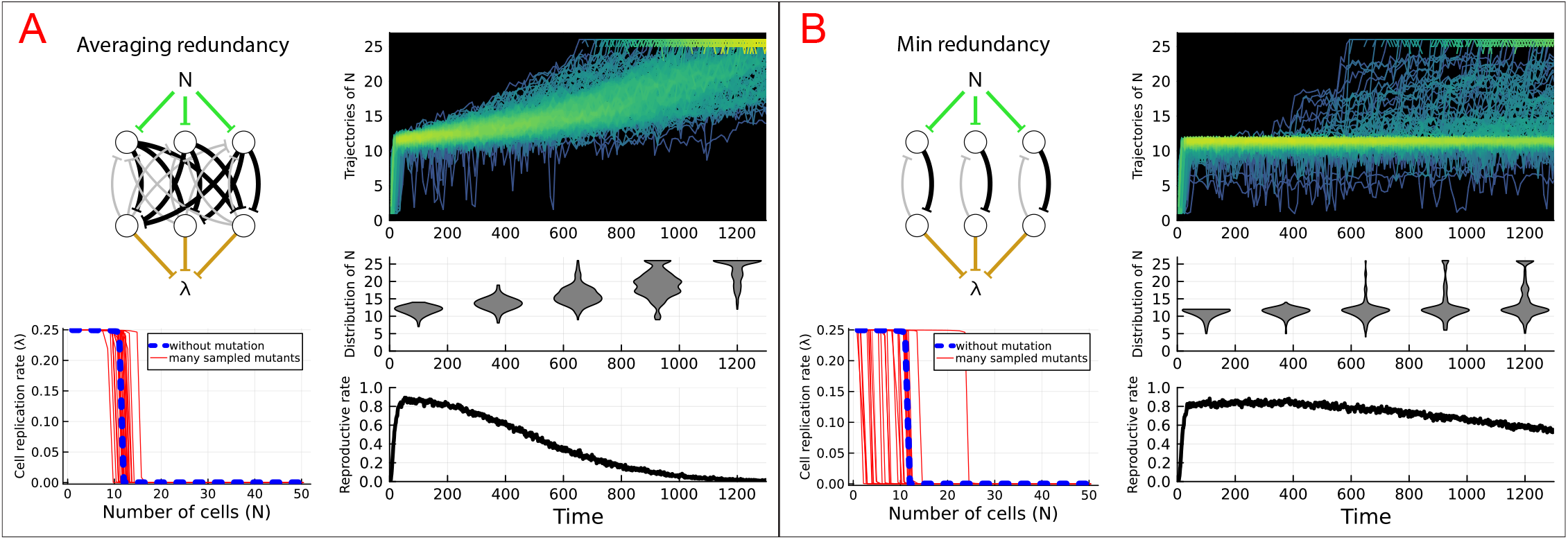
Response to mutation for averaging redundancy vs. min redundancy of the simple step motif. (A, top left) An example network structure is shown formed from three redundant simple step motifs. (A, bottom left) The distribution of many mutant cell replication functions are plotted alongside the nonmutant replication function, where mutant networks were created by independently sampling a Poisson distributed number of mutations to occur at a rate proportional to the number of edges in the network. (A, top right) 200 parsimonious creatures were independently simulated without organism death, where trajectories of *N* over time in each parsimonious creature. Overlapping trajectories are plotted brighter to illustrate density. (A, center right) The distribution of *N* among all simulations of parsimonious creatures over time are shown as violin plots. (A, bottom right) The average reproductive rate of all simulations of parsimonious creatures is plotted over time. (B) Same tests performed as in A, but for min redundancy. For illustrative purposes, a 16 fold averaging redundancy network and a 4 fold min redundancy network were tested in simulations of parsimonious creatures shown in panels A and B respectively.

This duplication creates significantly more interactions in the network which average out the effect of a single mutation on the replication function (Fig. 3, S1) in both a duplicated feedforward motif and a duplicated simple step motif, slowing hyperplasia in simulations. We explore this property analytically in the supplemental material. Briefly, the rate of somatic mutations is increased because of the larger target presented by a larger GRN. However, compensating feedbacks in the network act to decrease the effective mutation magnitude, and these combined effects reduce the effect of mutations on the mutant replication function (Fig 3. We refer to this form of motif redundancy as averaging redundancy.

The analytic results described in the Supplement predict that as the amount of averaging redundancy approaches infinity, fitness loss due to hyperplasia will be asymptotically reclaimed. However, the results of our simulations found that fitness increases slowly with respect to averaging redundancy network size (Fig. 4). Averaging via motif duplication is a brute force approach which often requires a very large GRN to significantly slow somatic evolution.

**Figure 4:**
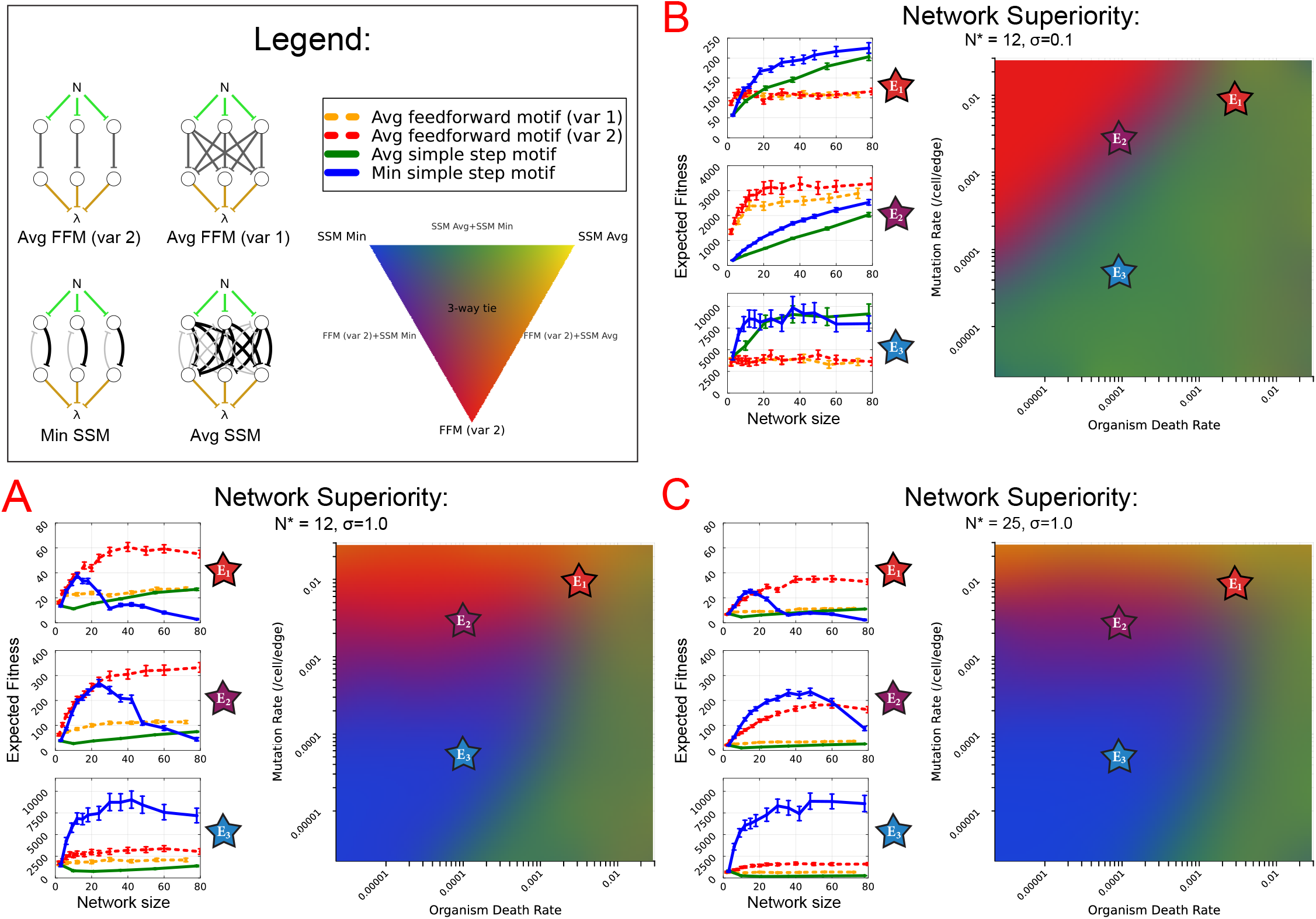
Fitness of multiple possible redundant network architectures formed from either the simple step motif or the feedforward motif are plotted with respect to network size in selected environments, while relative superiority is visualized in heatmaps for a grid of many possible environments. Example redundant network architectures formed from 3 repeated motifs are shown in legend. (A, left) Each plot focuses on a specific combination of organism death rate and mutation rate indicated by the stars *E*_1_, *E*_2_, and *E*_3_ (see text for details). To account for hedging, homogeneous 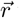 was optimized using Bayesian Optimization prior to estimating fitness of each network tested. Expected fitness was estimated by measuring fitness for many independent simulations of parsimonious creatures with the exact same optimized network, and 95 percent confidence intervals were estimated using bootstrapping. (A, right) A log-scaled grid of organism death rates and mutation rates is colored indicating probability that each of the following three network architectures has the highest fitness for any size up to the maximum network size (80 mutatable parameters) according to an RYB color palette: feedforward motif averaging redundancy variant 2 (red), simple step motif averaging redundancy (yellow), and min redundancy (blue). Feedforward motif averaging redundancy variant 1 is not included in heatmaps due to low relative fitness everywhere. To calculate network architecture superiority in heatmaps, 20 replicates of fitness were collected in simulations of parsimonious creatures for each position in a 12×127 grid of mutation rates and organism lifespans and across all network sizes independently for all network types, where 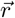 for each redundant network architecture and size was optimized for hedging utilizing a spatial evolutionary algorithm across the grid of environments. Bootstrapped t-tests were performed at each position in the grid of environments to calculate the probability that a particular network architecture has higher fitness than all other network types, then these three probabilities were colored at each position according to an RYB color gamut. Mutation magnitude is *σ* = 1.0 and *N*^*\**^ = 12. (B) Same tests as in A with mutation magnitude decreased from *σ* = 1.0 to *σ* = 0.1. (C) Same tests as in A with *N*^*\**^ increased from 12 to 25.

### Min redundancy

Suppose two redundant motifs detect signal and regulate the cell’s replication independently from one another. To limit hyperplasia, a sensible line of logic for a cell to follow could be “replicate according to the source of information that would regulate replication most conservatively.” Pleasingly, this can be achieved with two independent simple step motifs which each strongly and independently inhibit *λ* in parallel without mutual interactions as is found in averaging redundancy. We call this min redundancy (Fig. 3). This works due to the nature of step functions. Because replication is inhibited in an all or nothing manner, the interaction controlling growth 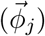 for each module can be arbitrarily strong, and when this is the case the relaxation of one SSM module will be negligible if the strong inhibition from the other module is still present. When mutant replication functions are sampled from a averaging redundancy network, mutant replication functions are symmetrically distributed, but if mutations are sampled in a min redundancy network, nearly all mutants have more conservative replication functions or have a neutral effect (Fig. 3). This forms a mechanism of epistasis [28], and furthermore creates increased fragility which Krakauer and Plotkin would describe as “antiredundancy” [29]. The philosophy of “taking the most conservative of multiple sources of information” seems to only be effective with step functions, as fully independent redundant copies of the feedforward motif is instead a variant averaging redundancy network (labeled feedforward avg var. 2 in data).

To clearly describe the behavior of a min redundancy network, we differentiate between *η*_*m*_, the threshold of an individual SSM module within the min redundancy network and *η*_*λ*_, the resulting threshold *N* of the cell’s replication function from the contributions of all SSM modules. For example, the resulting *λ* in 3 fold min redundancy network can be abbreviated as *η*_*λ*_ = min(*η*_1_, *η*_2_, *η*_3_). Therefore, a cell lineage within a parsimonious creature with min redundancy only suffers a more aggressive replication function if that lineage mutates to have a larger *η*_*m*_ in every one of its SSM modules. Any mutation which decreases the lowest *η*_*m*_ among all SSM modules will decrease *η*_*λ*_. If *η*_*λ*_ *< N*^*\**^, the cell will be selected against relative to nonmutants. This helps the organism remain robust to somatic evolution: hyperplasia is absent until neutral mutations that increase *η*_*m*_ in each of the SSM modules accumulate.

However, we must add one slight caveat. When min redundancy is used, mutant lineages which have accumulated more of these neutral mutations will have a small selective advantage when we consider further mutation. Consider a cell from a parsimonious creature with a min redundancy network in which *η*_*m*_ in one of its SSM modules has already mutated to have *η*_*m*_ much larger than *η*_*λ*_. If this cell receives another mutation in the same SSM module that then decreases *η*_*m*_, *η*_*m*_ is still likely to be greater than *η*_*λ*_, and the overall effect of this mutation which decreased *η*_*m*_ is in fact neutral. In contrast, in an unmutated min redundancy network with uniform *η*_*m*_ across all SSM modules the same mutation would have lowered *η*_*λ*_, resulting in the cell being selected against. Therefore, when min redundancy is used, the selective pressure against mutants will decrease as a mutant cell lineage accumulates neutral mutations, and emergence of hyperplasia will accelerate as neutral mutations accumulate.

### Life history parameters determine optimal redundant network architecture

We have described 2 motifs which can be used to regulate N in parsimonious creatures, as well as 2 forms of redundancy by which these motifs may be duplicated. However, we need to quantify whether the feedforward motif, the simple step motif, as well as whether to duplicate either of these motifs to form averaging redundancy or min redundancy will optimize fitness of parsimonious creatures, and in which environments a particular network architecture will be favorable. Furthermore, this may also depend on the amount of times a motif may be be duplicated. To test this, we considered 4 network architectures: Averaging redundancy of the simple step motif, min redundancy of the simple step motif, and the two variant forms of averaging redundancy of the feedforward motif. The amount of redundancy tested ranges from a single motif to the amount of redundancy that forms a GRN of 80 mutateable edges. We chose three environments to measure expected fitness of each network architecture with respect to network size.

- *E*_1_: lifetimes are short and mutation rate is very high,
- *E*_3_, lifespans are long and mutation rate is low,
- *E*_2_: intermediate environment between *E*_1_ and *E*_3_.

We set a baseline for two additional parameters of *N*^*\**^ = 12 and *σ* = 1.0 (Fig. 4A).

In the baseline case, feedforward motif averaging redundancy (var 2) either matches or exceeds fitness of all other network architectures in environments *E*_1_ and *E*_2_. We observe in *E*_3_ that min redundancy is superior at all network sizes only when mutation rate and organism death rate are sufficiently low. Furthermore, fitness peaks at intermediate network sizes for min redundancy in *E*_1_ and *E*_2_. We observe similar results when *N*^*\**^ is increased, where all network types decrease in fitness, but this decrease is least drastic for min redundancy (plots in Figure 4C). Furthermore, when *σ* decreases, we observe a significant reordering of network superiority in *E*_1_ and *E*_2_, while networks which modularize the same motif have approximately equal fitnesses in *E*_3_ at most network sizes (plots in Figure 4B). The latter of these observations is due to the significantly decreased overall mutational load, leading to rapid increases in fitness to the maximal fitness asymptote with respect to network size.

A more comprehensive investigation of parameter space reveals discrete territories across a grid of mutation rates and organism lifespans where each network architecture is superior given any network size up to 80 mutateable parameters (heatmaps in Fig. 4). When mutation rate and/or organism lifespan are sufficiently small, somatic mutations have little effect on a parsimonious creature’s fitness, and all network types which form a step-like replication function have equally high fitness as they are equally unperturbed by somatic evolution in this environment. When organism lifespan is long, but mutation rate is very low, min redundancy is superior due to the ability to delay hyperplasia until neutral mutations accumulate. However, when mutation rate is high, the inherent robustness of the feedforward motif is superior. Notably, we failed to find an environment where SSM averaging redundancy is superior to all other network architectures below a network size of 80 mutateable parameters, though we expect this network architecture to eventually dominate as network size approaches infinity.

When mutation magnitude is lowered from *σ* = 1.0 to *σ* = 0.1, we see a drastic reduction in the zone where min redundancy is superior, instead having equal fitness to SSM averaging redundancy. Moreover, this shrinkage increases with larger maximum network size (Fig. S3). Lowering *σ* makes each mutation less potent, until a sudden transition where feedforward motif networks are favored. This indicates that if mutation magnitude is small, the choice between min redundancy and averaging redundancy is of minimal consequence, both methods reach the maximum fitness asymptote quickly with respect to network size in most environments.

### Increasing adult cell count (*N*^*\**^) decreases fitness of averaging redundancy, but less so for min redundancy

If we increase from *N*^*\**^ = 12 to *N*^*\**^ = 25, we see an expansion of territories where min redundancy is superior (Fig. 4) However, when we plot fitness in selected environments, we observe general lower fitness due to accelerated hyperplasia at earlier timescales (Fig. 4). Moreover, this decrease in fitness is not largely attributable to longer times to reach *N*^*\**^, as a general decrease in fitness is still observed when simulations are configured such that organisms do not die until *N* reaches *N*^*\**^ (Fig. S5).

In the case of averaging redundancy, mutations always have an effect, even if small, and will be selected for at greater resolution and strength, while drift will be weaker at higher cell population sizes, significantly accelerating hyperplasia. With min redundancy, the decreased fitness at higher *N*^*\**^ is small compared to the decrease seen in other network types (Fig. 4). When more cells are in competition, the forces of selection on cells will be stronger, which results in two relatively weak processes that decrease fitness of min redundancy at higher *N*^*\**^. First, there is a small selective advantage of “neutral” mutations in min redundancy networks which will eventually lead to hyperplasia and will be selected at higher strengths for higher *N*^*\**^. Second, the rate of stochastic tunneling is greater at larger population sizes [30] and thus the accumulation of all neutral mutations in a min redundancy network for a cell to become hyperplastic can occur at earlier timescales.

### Large min redundancy networks are prone to ‘paralysis’

Large min redundancy networks often have decreased fitness, with fitness optimized at a moderate number of SSM modules in *E*_1_ and *E*_2_ (Fig. 4). At first glance, this result appears counterintuitive, as adding more SSM modules in min redundancy should increase the number of neutral mutations required to achieve hyperplasia. However, increasing network size creates a larger target, leading to a proportional increase in net mutation rate, of which a single mutation is very likely to lead to hypoplasia. We refer to this phenomenon as “bureaucratic paralysis”.

The first driver of bureaucratic paralysis is senescent collapse, which occurs when net mutation rate is sufficiently large that hypoplastic mutants (mutants with replication functions less aggressive than the nonmutant type) arise faster then they can be replaced by nonhypoplastic cells. Stated another way, selection will no longer be strong enough to maintain a quasispecies if the genome becomes very large and fragile [31, 32]. Consider an idealized example.

Suppose that extreme hypoplastic mutants arise which neither divide and nor back mutate to being nonhypoplastic. We can classify all cells as either nonhypoplastic cells, or nonreplicating hypoplastic cells. If these nonreplicating mutants arise often, nonhypoplastic cells must constantly replicate to compensate. Let *Q* be the number of nonhypoplastic cells and *µ*_*hypo*_ be the rate of formation of hypoplastic mutant cells. The dynamics can be simply described as

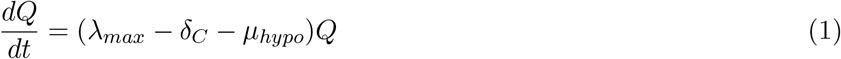

Where *λ*_*max*_ is the maximal cell replication rate and *δ*_*C*_ is the constant cell death rate.

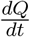 will be negative when *µ*_*hypo*_ exceeds *λ*_*max*_ *− δ*_*C*_, at which point the number of nonhypoplastic cells *Q* will on average decrease over time until the organism fully senesces at N=1 cells (Fig. 5).

**Figure 5:**
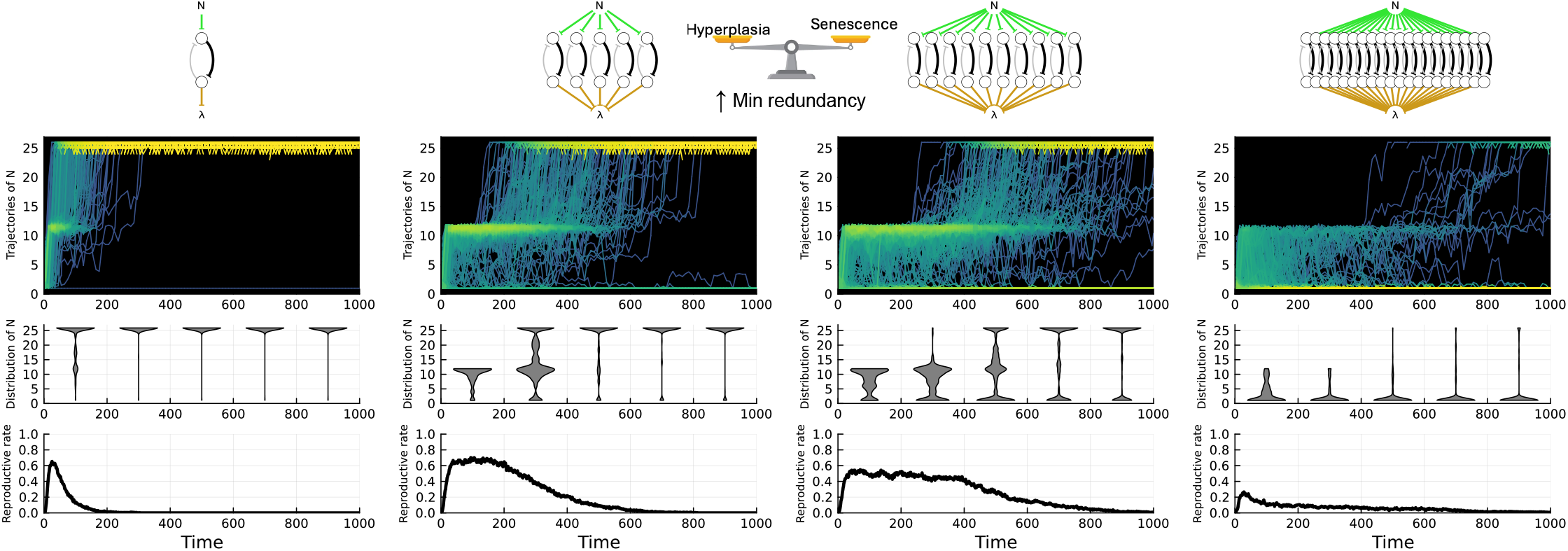
Simulations of parsimonious creatures with increasing min redundancy from left to right. Network topologies shown indicate the amount of min redundancy tested in the below plots. 200 parsimonious creatures were independently simulated with the simple step motif in the absense of organism death, where trajectories of *N* are plotted over time for each parsimonious creature. Overlapping trajectories are plotted brighter to illustrate density. Bright yellow bands at *N* = 25 and/or *N* = 1 are present due to *N* = 25 being an absorbing state, and organisms at *N* = 1 cannot undergo cell death. The distribution of *N* among all simulations of parsimonious creatures over time are shown as middle violin plots. Bottom plots show the average reproductive rate of all simulations of parsimonious creatures over time.

However, this idealized analysis ignores drift created by stochastic cell replication and cell death. Due to the fact that somatic cells are asexual, drift will drive Muller’s ratchet, ensuring that hypoplastic mutants will eventually replace all nonhypoplastic mutants given enough time in a finite somatic cell population, even when 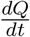 is positive [33]. The two forces of senescent collapse and Muller’s ratchet operate differently: the first is strong and immediate when *µ*_*hypo*_ exceeds *λ*_*max*_ *− δ*_*C*_, while the latter operates more gradually dependent on *µ*_*hypo*_ and *N*^*\**^.

In summary, bureaucratic paralysis occurs when an increase in network size increases the rate and proportion of hypoplastic mutants sufficiently to decrease expected lifetime fitness. Thus high min redundancy will increase the frequency and proportion of hypoplastic mutant formation, accelerating Muller’s ratchet and potentially crossing a threshold to cause senescent collapse. These two forces will decrease the force of selection maintaining a population of nonhypoplastic cells such that the organism senesces. This acts to disfavor large min redundancy networks, and this effect is accelerated by a higher mutation rate.

## Discussion

How can cells build gene regulatory networks that effectively regulate cell number in the face of somatic mutations? We have designed the parsimonious creature, an in-silico multicellular organism which gains fitness only when it possesses the right number of cells, and thus must mediate against the hyperplasia that results from somatic mutations that grow too quickly and outcompete unmutated cells.

### Summary of key results

Using the framework of our model of parsimonious creatures, we discovered two minimal networks, each with two nodes, that can respond to current cell number with decreased replication. The feedforward motif regulates cell growth only weakly, leading to low fitness in the absence of mutation, but is robust when mutations are introduced. The simple step motif has a strong negative feedback that regulates cell number very tightly around the optimal size, and thus has high fitness in the absence of mutation, but is much less robust to mutation.

By using these motifs as building blocks to form larger redundant networks, fitness can be maintained for longer periods in the face of somatic evolution despite the larger mutational target presented by the larger network. The robust GRN architectures described here do not rely on the direct detection of DNA damage, such as the famous “guardian of the genome” p53, but rather act via the altered GRN dynamics after mutations have already occurred.

We investigate two opposing strategies: averaging redundancy, which takes the average of multiple sources of information, and min redundancy, which adopts the most conservative source. Averaging redundancy can be formed by fully duplicating a functional motif, and analytic solutions suggest it effectively acts to reduce the magnitude of the effect of mutations on cell replication for both the feedforward motif and simple step motif. Averaging redundancy networks asymptotically reclaim all fitness lost to somatic mutation as network size increases (Fig. 4). However, these trends can be very slow with respect to network size, and transiently decrease fitness at moderate sizes when mutation magnitude is high.

Multiple redundant independent simple step motif modules form min redundancy, which adopts the minimum threshold *N* of all simple step motifs in the network. This phenomena is restricted to the simple step motif due to the all or nothing nature of step functions. Min redundancy asymmetrically separates mutations into two categories: those that either cause hypoplasia (reduced cell number), or are neutral to the cell’s overall replication function (Fig. 3). Hyperplasia is therefore delayed until all neutral mutations have accumulated when min redundancy is used.

However, parsimonious creatures that employ large min redundancy networks suffer from what we call “bureaucratic paralysis,” a counterintuitive decrease in fitness after increased redundancy is added to a network. Because net mutation rate is proportional to network size, hypoplastic mutants generate at higher frequency with higher min redundancy, and furthermore the probability of hypoplastic mutant reverting to a nonhypoplastic one becomes very low. From this follows the two component forces of bureaucratic paralysis: Muller’s ratchet, where hypoplastic mutants take over the population due to drift, and senescent collapse, where the rate of hypoplastic mutant formation exceeds the rate that nonhypoplastic cells can replace them. Overall, while high min redundancy constrains the paths to hyperplasia, it also vastly expands the paths to hypoplasia such that selection can no longer maintain a stable cell population.

### Could these forms of motif redundancy be used by multicellular life?

The evolutionary genomic formation of large averaging redundancy networks is fairly straightforward and could naturally arise from duplications, but forming a high min redundancy networks requires many cycles of duplication followed by the severing of interactions between motifs. During this process, min-avg hybrid networks become likely to form (Fig. S6A). Our testing of intermediate min-avg hybrid networks indicates that the evolutionary path is likely to be directed toward relatively pure min redundancy networks rather than min-avg hybrids (Fig. S6, S7). We have shown that large min redundancy networks can display bureaucratic paralysis, putting a cost on network size independent of direct costs, such as metabolic costs. We believe these limitations will constrain the search for multicellular gene regulatory networks robust to somatic evolution in multicellular organisms. We make the following prediction: relatively small and pure min redundancy networks will be found in the life history environments in which min redundancy is favorable: mutation magnitude is large, somatic cell pools are maintained competitively, and mutation rate is significant, but not overwhelming. Otherwise, we expect large averaging redundancy to be found.

### Stem cell dynamics could determine which form of redundancy is favored

We found that at higher cell population sizes (*N*^*\**^), all network types decrease in fitness, though the decrease is comparatively small for min redundancy than for averaging redundancy, leading to an expansion of environments where min redundancy networks are superior (Fig. 4A, 4B). At higher cell population sizes, the force of selection acts strongly to increase the rate of hyperplastic mutant takeover in parsimonious creatures with averaging redundancy, significantly slowing the rate at which they reclaim fitness with respect to network size. Meanwhile, if min redundancy is used, the acceleration of hyperplasia solely occurs via stochastic tunneling, and the increased force of selection instead acts to strengthen the removal of mutant cells inherent to min redundancy.

We do not know if these trends will hold at physiologically relevant somatic cell populations that often number in the millions in multicellular lifeforms. However, space and the locality of stem cell competition can greatly reduce these numbers. For example, in mice each intestinal crypt is believed to be a “closed niche” maintained by 6-7 stem cells [34]. A more “open niche” is supported in bone marrow, where thousands or millions of stem cells may be in competition due to routine long distance migrations [35]. Our results suggest that these differences in the number of stem cells in competition in different contexts could encourage the deployment of fundamentally differing forms of redundant GRN structures not only between different species but also between different cell types.

While the bulk of our results have focused on mutations solely occurring in somatic cells with a fully sequestered and nonmutating germline, Fig. S2 suggests that the ranking of network architectures can significantly change when germline mutations are present. This data suggests that in an evolutionary model over many generations of parsimonious creatures, averaging redundancy would become favored due the accumulation of mutations in the germline that disfavor min redundancy. However, this conclusion ignores the possibility of germline cell competition, which appears to vary significantly among species. For example, an open niche model of spermatogonal stem cells is supported in mouse [36, 37], but not in comparatively short lived Drosophila and C. *elegans* [38].

### Lessons for synthetic biologists

Throughout this work we have designed GRNs to be robust to somatic evolution and have tested them in parsimonious creatures, free of constraints governing whether these GRNs can form via stepwise evolution, or direct metabolic costs. Despite this freedom, we discover fundamental tradeoffs governing a redundant GRNs robustness to somatic evolution. We believe these results could be particularly noteworthy for synthetic biologists. For instance, in yeast cultures engineered to produce insulin, “cheater cells” which mutate to produce less insulin will then have more resources to devote to reproduction and will be at a selective advantage, eventually lowering the yield of cell factories [39].

The problem of divergence in cell populations have real monetary costs that synthetic biologists would be interested in reducing, and we offer the following guidance when designing a gene regulatory network robust to it: First, there is a fundamental decision whether to lower the effective mutation magnitude using averaging redundancy, or neutralize hyperplastic mutations via min redundancy until neutral mutations accumulate. Nevertheless, even very high averaging redundancy appears to become very ineffective as the number of cells in competition increases. Furthermore, if a mutant removing strategy is chosen, bureaucratic paralysis is a limitation which must be considered, as we have seen in large min redundancy networks. Therefore, we would suggest testing a model of the expected distribution of mutants in their designed gene regulatory network combined with the specific dynamics of cell replacement in their desired system to accurately determine which form of redundancy, as well as what amount of redundancy, is optimal to reduce in-vivo testing.

## Methods

**Table 1:**
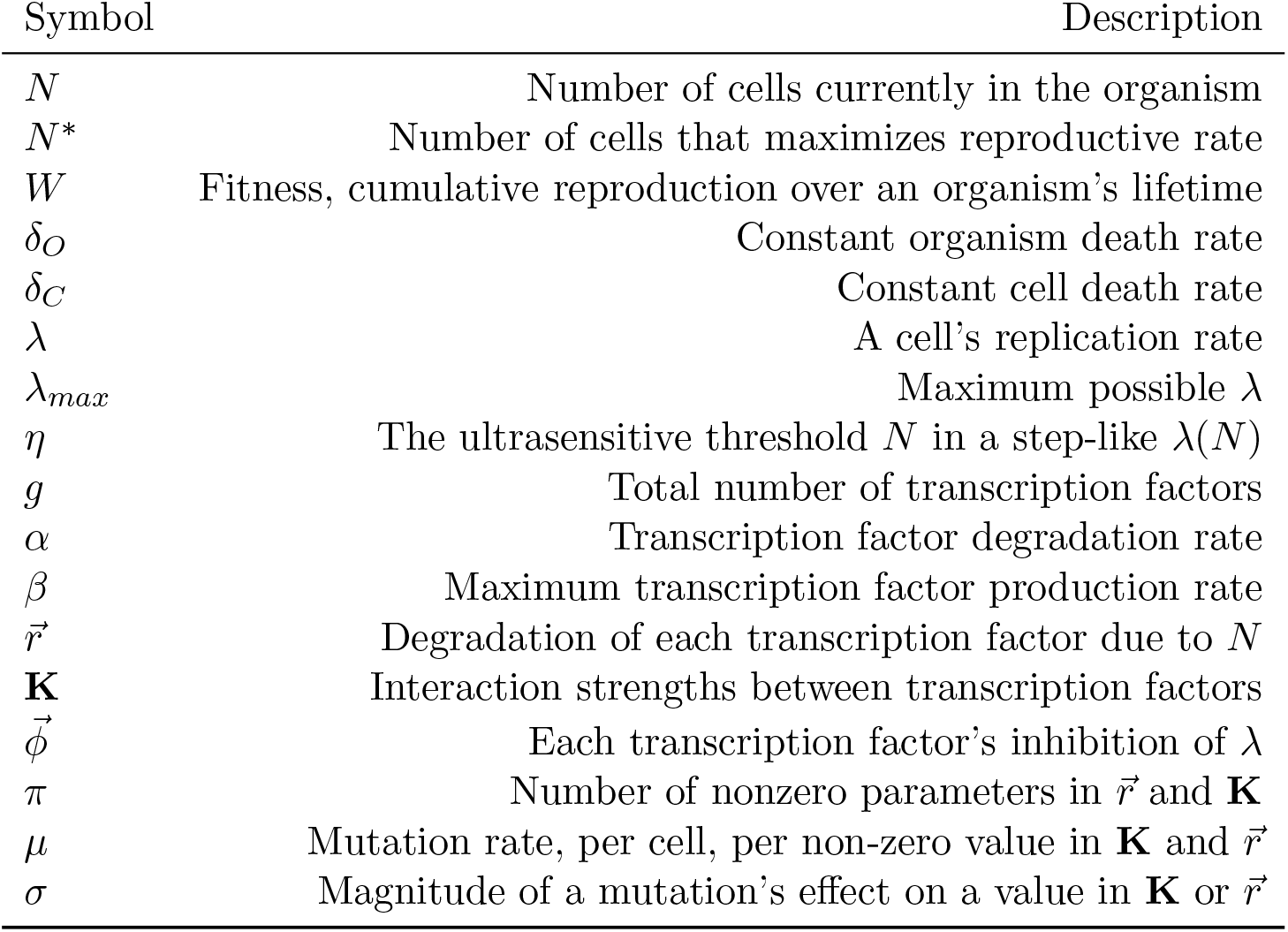
Brief description of symbols.

| Symbol | Description |
|---|---|
| *N* | Number of cells currently in the organism |
| *N*<sup>\*</sup> | Number of cells that maximizes reproductive rate |
| *W* | Fitness, cumulative reproduction over an organism’s lifetime |
| *δ*<sub>*O*</sub> | Constant organism death rate |
| *δ*<sub>*C*</sub> | Constant cell death rate |
| *λ* | A cell’s replication rate |
| *λ*<sub>*max*</sub> | Maximum possible *λ* |
| *η* | The ultrasensitive threshold *N* in a step-like *λ*(*N*) |
| *g* | Total number of transcription factors |
| *α* | Transcription factor degradation rate |
| *β* | Maximum transcription factor production rate |
| $\vec{r}$ | Degradation of each transcription factor due to *N* |
| **K** | Interaction strengths between transcription factors |
| $\vec{\phi}$ | Each transcription factor’s inhibition of *λ* |
| *π* | Number of nonzero parameters in $\vec{r}$ and **K** |
| *μ* | Mutation rate, per cell, per non-zero value in **K** and $\vec{r}$ |
| *σ* | Magnitude of a mutation’s effect on a value in **K** or $\vec{r}$ |

### Modeling parsimonious creatures

A parsimonious creature begins life from a single somatic cell that serves as the progenitor of the entire organism. This first somatic cell and its descendant cells can divide or die, creating an organism of size *N* measured as the current number of cells. We assume that the germline and the soma are segregated, but we do not explicitly include germline evolution, and parsimonious creatures are born with a predefined genome rather than one that is inherited by another organism. Nevertheless, we still measure “reproductive rate” as a measurement of how successful a parsimonious creature would be if reproduction were determined by W(N), a Gaussian function which is maximized at *N* = *N*^*\**^. This favors tightly regulated cell replication that keeps *N* as close to *N*^*\**^ as possible over time. We assume a constant background organism death rate *δ*_*O*_ and cell death rate *δ*_*C*_. Lifetime fitness will be maximized for organisms who replicate the soma quickly to size *N*^*\**^ and inhibit further replication unless *N* drops below *N*^*\**^ due to cell death.

The genome within each cell encodes a set of transcription factors (TF). For simplicity and to reduce the probability of oscillatory solutions, we assume that all interactions are inhibitory, with a Michaelis-Menten form [40]. Thus, each transcription factor species is expressed constitutively unless inhibited by signal as described below, or inhibited by one or more transcription factors, possibly including itself, via repression of a transcription factor protein competitively binding to the promotor. We assume that transcription and translation are coupled.

For example, the concentration *T* of a single TF that represses itself via promoter competition obeys

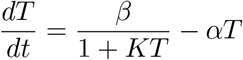

Here *β* and *α* represent the transcription and degradation rates, and *K* is the association constant of the transcription factor to its own promoter.

We generalize this to a system of TFs, each capable of regulating or being regulated by all other transcription factors, including self-interactions. Assuming the number of unbound transcription factors is much greater than the number of DNA binding sites, the concentration *T*_*i*_ of transcription factor *i* obeys

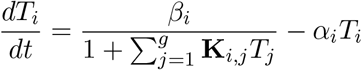

Where *g* is the number of TF genes in the system, **K** is the matrix of interactions between transcription factor species, which describes a directed weighted graph. Transcription factor *j* reduces the production rate of transcription factor *i* with parameter *K*_*ij*_, with the overall topology of the network defined by the set of nonzero values in **K**.

These transcription factor dynamics are linked to organism size in two ways. First, any subset of these transcription factors may respond to the current number of cells in the organism through an increased decay rate linearly proportional to *N*. Although measuring the number of cells is in reality a complex task fraught with uncertainty, we assume *N* is a pure signal that exactly measures the current number cells in the organism, modifying TF dynamics to

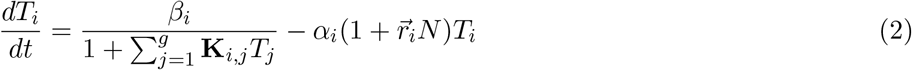

The vector 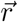 describes each TF’s sensitivity to *N*.

Second, any subset of transcription factors may also directly inhibit cell replication rate *λ*.

For any *N*, we assume that the GRN in each cell reaches equilibrium more quickly than changes in *N* due to replication or cell death, and denote the equilibrium concentration of transcription factor *i* as *T*_*i*_ for convenience. The equilibrium concentrations of transcription factors controls the rate of cell replication, modeled as the expression of a single non-transcription factor output gene, *λ*:

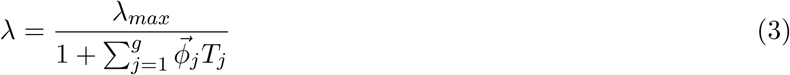

The vector 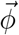 describes how strongly each TF represses cell replication rate *λ* below the maximum of *λ*_*max*_.

When a cell divides, its two daughter cells inherit **K**, 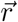 and 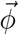 which may have mutated at any point to alter one or more of the parameters to produce a perturbed replication function. As cells in organism replicate, a mosaic of cells will form from different lineages of somatic mutants which may divide at different rates as a function of *N*.

### Mutations

We assume that each nonzero value in 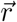 or **K** can mutate independently at constant rate *µ* by being multiplied by a random value drawn from a lognormal distribution with a a log mean of 0 and log standard deviation *σ* controlling the mutation magnitude. Mutations are thus log-symmetric and equally capable of increasing or decreasing interaction strengths. The net mutation rate per cell will be proportional to the number of nonzero values in 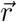 and **K**. Because mutations to 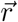 and **K** are upstream of components which can compensate for the perturbation, little can be done to respond to mutations in exclusively downstream 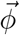. Therefore, we have chosen to exclude somatic mutations to 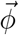.

### Simulation based model of parsimonious creatures

In the stochastic simulation based model of parsimonious creatures, cell replication, cell death, mutation, and organism death occur at rates *λ*(*N*), *δ*_*C*_, *µ*, and *δ*_*O*_ respectively. These are modeled as instantaneous events in continuous time using the Gillespie algorithm [41]. With our separation of timescales assumption, when a network parameter changes through mutation, or input signal *N* changes, the concentration of transcription factors instantly reaches a new equilibrium. To implement this, we analytically or numerically solve the system for equilibrium in each cell to calculate the replication rate of each cell following each event. Fitness accumulates over each time step *dt* as *∑W* (*N*)*dt*, where fitness of the organism is determined either at organism death, or if *N >* 50.

### Deterministic modeling of parsimonious creatures in the absence of mutation

A deterministic analog to a Gillespie algorithm is a system of master equations, which model an infinite population of organisms which may transition at some rate to an alternative state with a different number of cells. The system is then solvable for equilibrium probability distribution of each N. This approach is computable only in a relatively small state space, which is the case if we assume mutation rate is zero. Organisms in an *N* cell state gain one cell through replication at rate *λ*(*N*) to transition to the *N* + 1 cell state or lose one cell through cell death at rate *Nδ*_*C*_, and each organism can die at rate *δ*_*O*_. Organisms are born at rate *δ*_*O*_, entering the system at the one cell state. As boundary conditions, single celled organisms cannot undergo cell death and we choose a maximum *N*_*max*_ = 50 to be purely absorbing. By solving for *λ*(*N*) separately as a look up table for each integer *N* between 1 and *N*_*max*_ for a given GRN, this can be written as a system of linear ordinary differential equations for the probability of residing in each state:

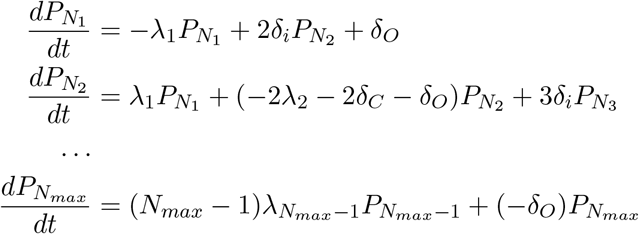

The vector *P*_*N*_ can be solved as equilibrium probabilities of an organism being at any number of cells. Expected fitness is

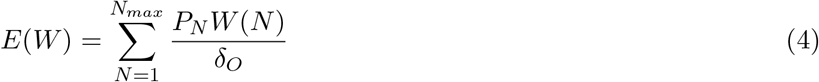

### Optimization

An organism that spends time *τ*_*N*_ with *N* cells has a cumulative lifetime reproduction (fitness), calculated as 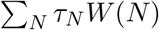, where *W* (*N*) describes reproductive rate as a function of cell number. In the absence of mutation (*µ* = 0), **K**, 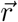, and 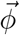 can be optimized to find a GRN that maximizes fitness using the purely nonmutant deterministic model of parsimonious creatures (Eq. (4)) to keep *N* constrained, for which we used blackbox optimization [12]. However, when *µ >* 0, homogeneous 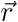 must be optimized for hedging. Due to the stochastic nature of simulations we used Bayesian optimization on log-transformed fitnesses [42]. In heatmaps of network superiority, a spatial evolutionary algorithm was employed across the grid to optimize homogeneous 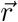 for hedging.

### Comparison of network types and sizes in selected environments

In selected environments *E*_1_ *− E*_3_, where each environment is a unique combination of base mutation rate and organism death rate, several sizes of min redundancy networks and averaging redundancy networks were tested which modularize either the feedforward motif or the simple step motif. This data was generated independently of data for heatmaps of network superiority. In each environment for each network size, *r* was chosen to be optimized for hedging to maximize fitness using bayesian optimization on log transformed fitnesses for normality. Independent replicate simulations are performed with this fixed *r* to collect final fitness values for a network optimized for hedging. Bootstrapping with 30000 replicates was then performed for these optimized *r* fitnesses to extract error bars.

### Heatmaps of network superiority

We comprehensively collected the fitnesses of different network types over a 127×127 grid of mutation rates and organism lifespans, optimizing *r* at each position on the grid independently for each network type and size. For data visualization, we capped maximum network size at either small networks (10 parameters), medium networks (30 mutatable parameters), or large networks (80 mutatable parameters), and chose the network size that maximizes fitness below each of these thresholds for each network type and in each position on the grid of environments. Using bootstrapping with a sampling of 30000, we then performed t-tests for each position in this grid of mutation rates and organism lifespans to quantify the probability that a particular network type has higher log mean fitness than all other network types. Due to the resultant noise across this grid, we applied a gaussian blurring kernel with a standard deviation of 10, and the resultant 3D vector of probability of each network superiority is then visualized for each position in the grid as a heatmap along an RYB color gamut.

## Data availability

All data was generated and plotted using the Julia programming language. Raw data to generate heatmaps are on [Zenodo](https://zenodo.org/records/23656540): 10.5281/zenodo.23656539. All other data and code, as well as docker images to generate this data and plots are available on GitHub: https://github.com/hyrumdiesen/ParsimoniousCreatures1

## Acknowledgments

We would like to thank Michael Werner for ideas on the framing of this work. Connor Shrader for editorial feedback. Jessica Brown, Richard Clark, Berten Earnshaw, James Gagnon, and Braxton Osting for guidance and comments in the early research phase. This work was supported in part by the 2025-2026 James E. Talmage Research Fellowship.

## Supplemental

### Algebraic solution and parameters for the simple step motif

Given the topology of interactions in the simple step motif, *T*_1_ inhibits *T*_2_ at association constant *k*_2_, and *T*_2_ inhibits *T*_1_ at association constant *k*_1_ via Michaelis-Menten dynamics, the number of cells in the organism, *N* acts a signal to increase the degradation of *T*_1_ at rate *r*, and finally *T*_2_ inhibits replication of the cell, *λ* at rate *ϕ*. Assuming transcription factor birth and decay rates are normalized to one, the system describing the simple step motif can be described as:

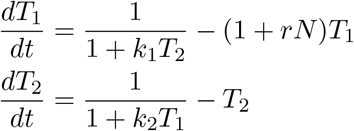

Due to the presence of a single feedback in the system, the dynamics of the simple step motif are algebraically solvable for equilibrium, assuming all parameters, variables and solutions are nonnegative and real numbers:

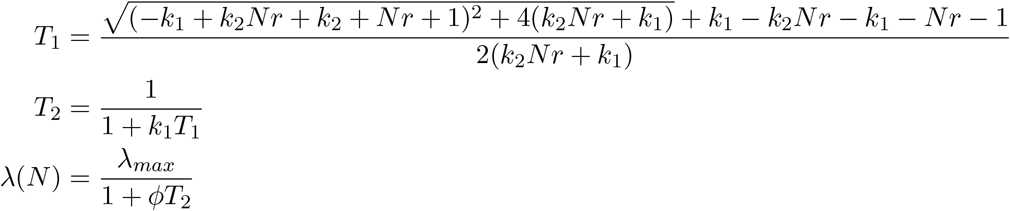

Blackbox optimization on our deterministic model of parsimonious creatures, which assumes *µ* = 0, optimized *r* = 0.174, *k*_1_ = 1.00 *\** 10^8^, *k*_2_ = 3.29 *\** 10^7^, *ϕ* = 2.31 *\** 10^4^. For simulations, *r* is optimized for hedging dependent on network size and life history parameters. For larger networks, 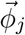 was normalized for the number of additional SSMs regulating *λ*.

### Hedging can partially recover fitness lost by somatic mutations

When somatic mutations occur in parsimonious creatures, a sensible strategy to reclaim long term fitness could be to “hedge” [43, 44] against somatic mutations by beginning life with with a nonmutant replication function that targets an *N* less than *N*^*\**^. We adopt the term “hedging” because a parsimonious creature could maximize fitness for an optimistic scenario in which it happens to never suffer somatic mutations for which we have optimized thus far. However, a parsimonious creature may be better off trading off fitness in the scenario in which no somatic mutations occur so that it may perform less terribly in the likely scenario in which it does suffer somatic mutations. For example, consider a parsimonious creature utilizing the simple step motif to maintain *N*^*\**^ = 12 cells to maximize fitness. When somatic mutations occur, an initial *η* = 12 is likely to be too optimistic for organisms that begin life at *N* = 1. By the time the organism reaches *N* = 6, *η* of the replication function of many cells could have already mutated from *η* = 12 to *η* = 24. Therefore, a reasonable network design might be to hedge initial nonmutant *η* to *η* = 6. Mutant cells might mutate from *η* = 6 to *η*_*m*_ = 12, creating an organism much closer the target size *N*^*\**^ = 12. Therefore, in the presence of somatic mutations, a parsimonious creature might “hedge” against somatic evolution by beginning life with a more conservative replication function. The amount of hedging required to maximize fitness will vary with expected lifespan and mutation rate, but is also dependent on the distribution of mutants due to network structure. To account for hedging, each network type and size in each environment was independently optimized with homogeneous *r* prior to estimating expected fitness.

### Averaging redundancy can be approximated as a single motif where mutation magnitude is decreased and mutation rate is increased

We found that mutating a single *K* parameter in an averaging redundancy network produces a nearly identical mutant replication function as the same network *K* with the geometric mean of all symmetric parameters after the mutation. This allows us to model all mutations as affecting a single value. Using this approximation for all mutations in an averaging redundancy network, we can analytically predict the effect of mutations in an averaging redundancy network as follows.

Given a single mutateable edge in **K** or 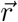 at any time, *t*, a Poisson distributed number of mutations has occurred on this parameter. Mutations multiply the value by a lognormally distributed number with mean 1, which when transformed in log space is an additive Gaussian random walk. The distribution of *Y* = log(*X*) at any time is a Poisson distributed mixture of Gaussians:

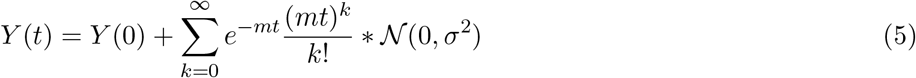

where *N* is the normal distribution, *Y* (0) is the initial log value, and *mt* is the expected number of mutations. Ross [45] finds a closed form solution for *Y* (*t*) with moments

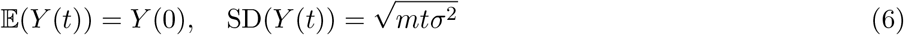

Where E and SD indicate the expectation and the standard deviation respectively. At time *t* with *π* mutateable parameters, we draw *π* independent samples from this distribution, exponentiate, and calculate the geometric mean. This predicts the following approximate equality of fitnesses in a single motif compared to averaging redundancy:

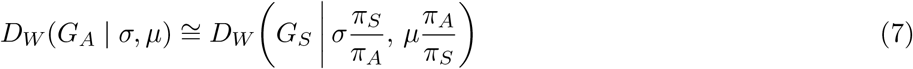

where *D*_*W*_ is the distribution of fitnesses, *G*_*S*_ is a GRN composed of a single module such as the feedforward motif or the simple step motif, *G*_*A*_ is an averaging redundancy network of multiple duplicated modules, and *π*_*S*_ and *π*_*A*_ are the number of mutating parameters in *G*_*S*_ and *G*_*A*_ respectively.

Finally, we tested an averaging redundancy network utilizing the simple step motif against a single simple step motif with adjusted mutation rate and mutation magnitude. We found very similar traces of N and average reproductive rate over time (Fig. S1), supporting our assertion that averaging redundancy can be approximated as a single motif where the mutation rate increases while mutation magnitude decreases.

### Moderate sized averaging redundancy networks dip in fitness under high mutation magnitude

A notable property of averaging redundancy is that fitness can transiently drop with a small to moderate network sizes, before increasing again for large averaging redundancy networks (Fig. 4). However, this effect is only visible in averaging networks which modularize the simple step motif (SSM modularizing), and if we decrease mutation magnitude (*σ*), the drop in fitness for moderate sized averaging redundancy networks disappears.

This observation supports the following explanation of this behavior: A mutation with a large mutation magnitude *σ* is likely to strongly translate the *η* of a step-like replication function, however, the translation of the *η* in hyperplastic mutant cells from *η* = 12 to *η* = 20 compared to *η* = 40 makes little difference from the perspective of the parsimonious creature who, given *N*^*\**^ = 12, effectively loses all fitness when *N >* 16. Therefore, if *σ* is already high when comparing a small averaging redundancy network to a single simple step motif, the increase in net mutation rate due to the higher number of parameters is not worth the decrease in effective *σ* at moderate averaging redundancy, culminating as a transient dip in fitness when mutation magnitude is large.

### Key constraints

We constrained the possible types of mutations in several ways. Although mutations in real cells may be biased toward loss-of-function, we chose symmetric mutations to discover topologies equally robust to mutations in both the “up” and “down” directions. In addition, for the bulk of our results we have assumed that each mutation affects only a single interaction between transcription factors, and that there is full independence between mutations. For example, nonsense mutations or deletions can knock out genes entirely with no backmutation. However, we do consider node mutations, where we see no significant reordering of network superiority.

Our second broad constraint governs how transcription factors are regulated by signal and by each other. We have assumed that all interactions are inhibitory, including signaling. We also assume that these inhibitory interactions all act through Michaelis-Menten competition for a binding site to inhibit transcription. Cooperativity and Hill dynamics are required to create a strong biological switch [46], and this could create a form of memory that could enhance robust GRN design.

### Node mutations

In our main data (Fig. 4), we have assumed that mutations affect one specific interaction (one value in **K** and 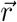) to perform “edge mutations”. Alternatively, a mutation could affect an entire node and change the strength of all of the inhibitory connections it creates by modifying the rate at which a transcription factor can be maximally produced, *α* to perform “node mutations”. We collected data for parsimonious creatures that experience node mutations rather than edge mutations in *E*_1_, *E*_2_, and *E*_3_ in the baseline case where *N*^*\**^ = 12 and *σ* = 1.0. First we must adjust for the fact that the network size is measured as the number of mutateable parameters, and is thus the number of nodes rather than the number of edges. For instance, large averaging redundancy networks now terminate at 12 nodes where they were no longer computationally feasible to simulate. Nevertheless, we do not observe major changes in network superiority compared to edge mutations.

### Germline mutations

In figure S2, we focus specifically on adding germline mutations in E1-E3, where a parsimonious creature has a chance of beginning life with mutations already present at N=1 at a germline mutation rate *γ* per mutatable edge in the GRN. Because these mutations are considered to be inherited from the germline and are present in the first cell at N=1, they will be inherited by all cells in the organism. We can see that dependent on the dose of *γ*, min redundancy begins to lose superiority to averaging redundancy, which is comparatively unaffected, and that the loss in fitness of high min redundancy to germline mutations increases with network size, appearing to accelerate bureaucratic paralysis.

### Hybrid min-avg networks

Through intermediate cycles of duplication and subfunctionalization, GRNs can be created that blend averaging and taking the minimum (Fig. S6A). For example, if we have a two-fold min redundancy, then duplicate one module but not the other.

In general, in a min-avg hybrid redundancy, the distribution of mutant replication functions has a smaller spread, similar to averaging redundancy, but take the minimum *η* of multiple averaged motifs. For example, a three-fold min redundancy network can be described with the logic: *η*_*λ*_ = min(*η*_1_, *η*_2_, *η*_3_), describing a min function with 3 arguments, and a three-fold averaging redundancy forms the logic: *η*_*λ*_ = avg(*η*_1_, *η*_2_, *η*_3_) a hybrid network nests averaging within a min statement, e.g *η*_*λ*_ = min(*η*_1_,avg(*η*_2_, *η*_3_)), and is a min function with 2 arguments, where the 2nd argument is averaged. In this simple min-avg hybrid redundancy, a mutation can be averaged within a min argument, but *η*_*λ*_ will ultimately take the minimum of an averaged *η* and a less averaged *η*. We adopt a shorthand notation to describe any min-avg hybrid redundancy, where for example *η*_*λ*_ = min(*η*_1_,avg(*η*_2_, *η*_3_)) is notated as the vector [1, 1, 0, 0, 0, 0], the network takes the minimum of two arguments, the first “averages” one ssm module, and the second averages two ssm modules.

We comprehensively tested all 31 pure min redundancy and pure averaging redundancy networks, as well as all 281 min-avg hybrid redundancy networks which can be created in parallel using 80 or fewer mutateable parameters in E1, E2, and E3 while optimizing homogeneous *r* for hedging. In all E1-E3, hybrid networks can increase fitness compared to pure min redundancy networks with the same number of min arguments (Fig. S7). To understand why, in Fig. S6 we subset the data to all networks which take the effective min() of 5 modules in E2. When mutation magnitude is small, we see that network size very strongly predicts expected fitness among hybrid networks, indicating that higher averaging redundancy increases fitness, regardless of the exact hybrid network type. However, as mutation magnitude increases, this trend breaks, indicating that decreases in fitness are merely due to the effect of moderate averaging decreasing fitness in parsimonious creatures as has seen previously in averaging redundancy networks at large mutation magnitude (Fig. 4): the mutation magnitude is large enough that an effective decrease in mutation magnitude in moderate averaging redundancy is not worth the accompanying increase in net mutation rate. When we label hybrid network type, we observe that hybrid networks which perform moderate amounts of averaging among any of it’s min() arguments has lower fitness, while pure min redundancy networks, or hybrid networks which combine small and large averaging among it’s min() arguments have higher expected fitness.

Assuming that duplication and subfunctionalization are the only actions available to form larger networks from the simple step motif, our data suggests implications for the evolvability of such networks. If mutation magnitude is small, min-avg hybrid networks may move relatively smoothly in the fitness landscape. However when mutation magnitude is large, low fitness barriers will exist for any min-avg hybrid which incorporates moderate averaging among any of it’s min arguments. For example, in an environment favorable to min redundancy, an even cycle of duplicate-subfunctionalize-duplicate-subfunctionalize etc, will require the lowest amount of compensatory evolution to form gene regulatory networks highly robust to somatic evolution, and may thus be restrained to pure or nearly pure min redundancy despite the fact that an asymmetric min-avg hybrid may return slightly higher fitness.

Overall, min-avg hybrid redundancy seem to predominantly have higher fitness than pure min redundancy only in environments where averaging redundancy performs well, and in particular when moderate averaging does not decrease fitness due to a large mutation magnitude.

**Figure S1:**
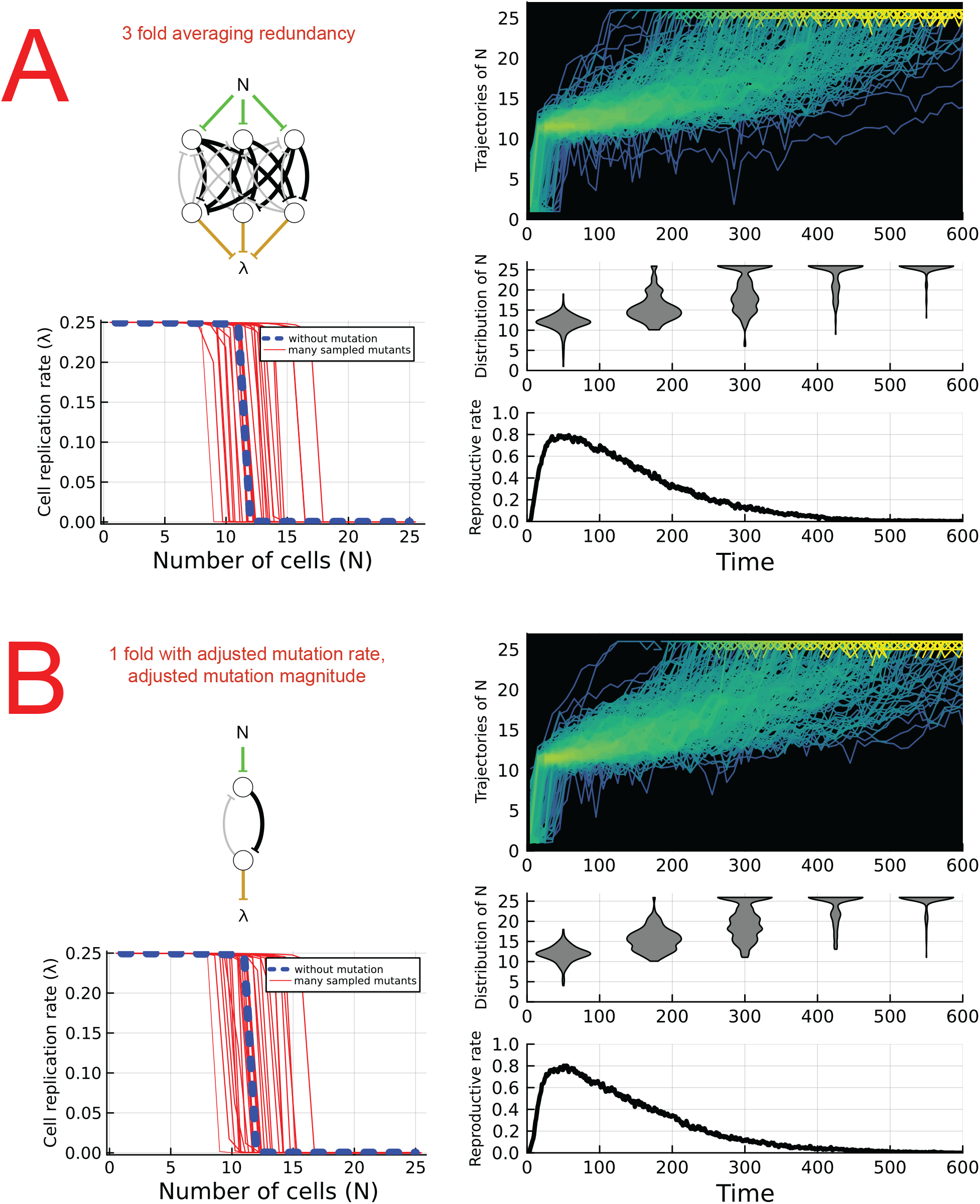
Response to mutation for averaging redundancy of the simple step motif vs. a single simple step motif with adjusted mutation rate and mutation magnitude. (A, top left) An example network structure is shown formed from three redundant simple step motifs. (A, bottom left) The distribution of many mutant cell replication functions are plotted alongside the nonmutant replication function, where mutant networks were created by independently sampling a Poisson distributed number of mutations to occur at a rate proportional to the number of edges in the network. (A, top right) 200 parsimonious creatures were independently simulated without organism death, where trajectories of *N* over time in each parsimonious creature. Overlapping trajectories are plotted brighter to illustrate density. (A, center right) The distribution of *N* among all simulations of parsimonious creatures over time are shown as violin plots. (A, bottom right) The average reproductive rate of all simulations of parsimonious creatures is plotted over time. (B) Same tests performed as in A, but for a single simple step motif with adjusted mutation rate and mutation magnitude (see equation 7 in supplement).

**Figure S2:**
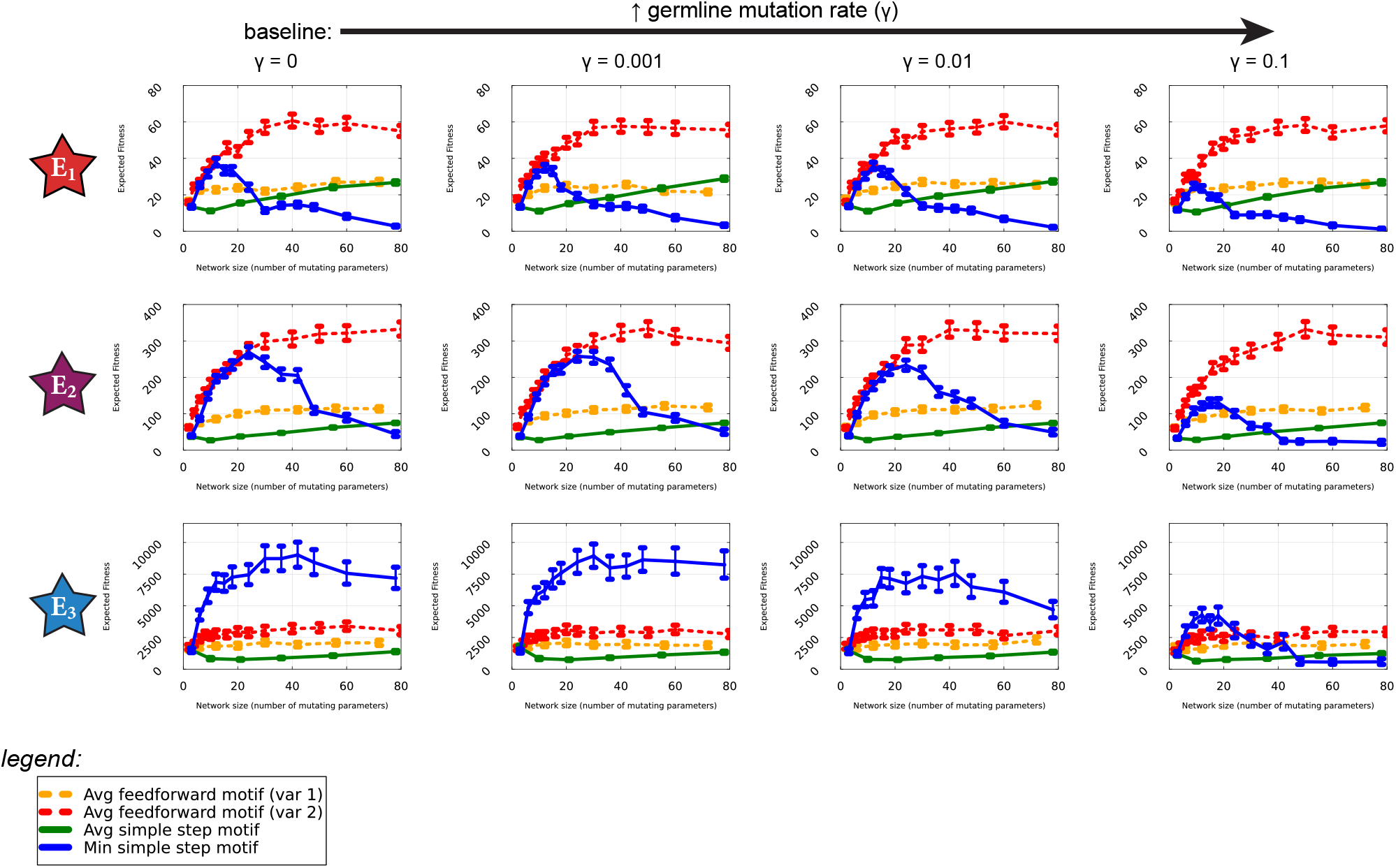
In each plot, expected fitness vs number of mutating parameters is plotted for feedforward averaging redundancy networks, ssm min redundancy and ssm averaging redundancy. Each plot focuses on a specific combination of organism death rate and mutation rate indicated by the stars *E*_1_, *E*_2_, and *E*_3_ (see text for details). The leftmost column of plots set the baseline life history parameters of *N*^*\**^ = 12 and *σ* = 1.0, while columns to the right indicate an increasing rate of germline mutations present in the initial cell of a parsimonious creature per cell per mutateable edge in a gene regulatory network. To account for hedging, homogeneous 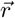 was optimized using Bayesian Optimization prior to estimating fitness of each network tested. Expected fitness was estimated by measuring fitness for many independent simulations of parsimonious creatures with the exact same optimized network, and 95 percent confidence intervals were estimated using bootstrapping.

**Figure S3:**
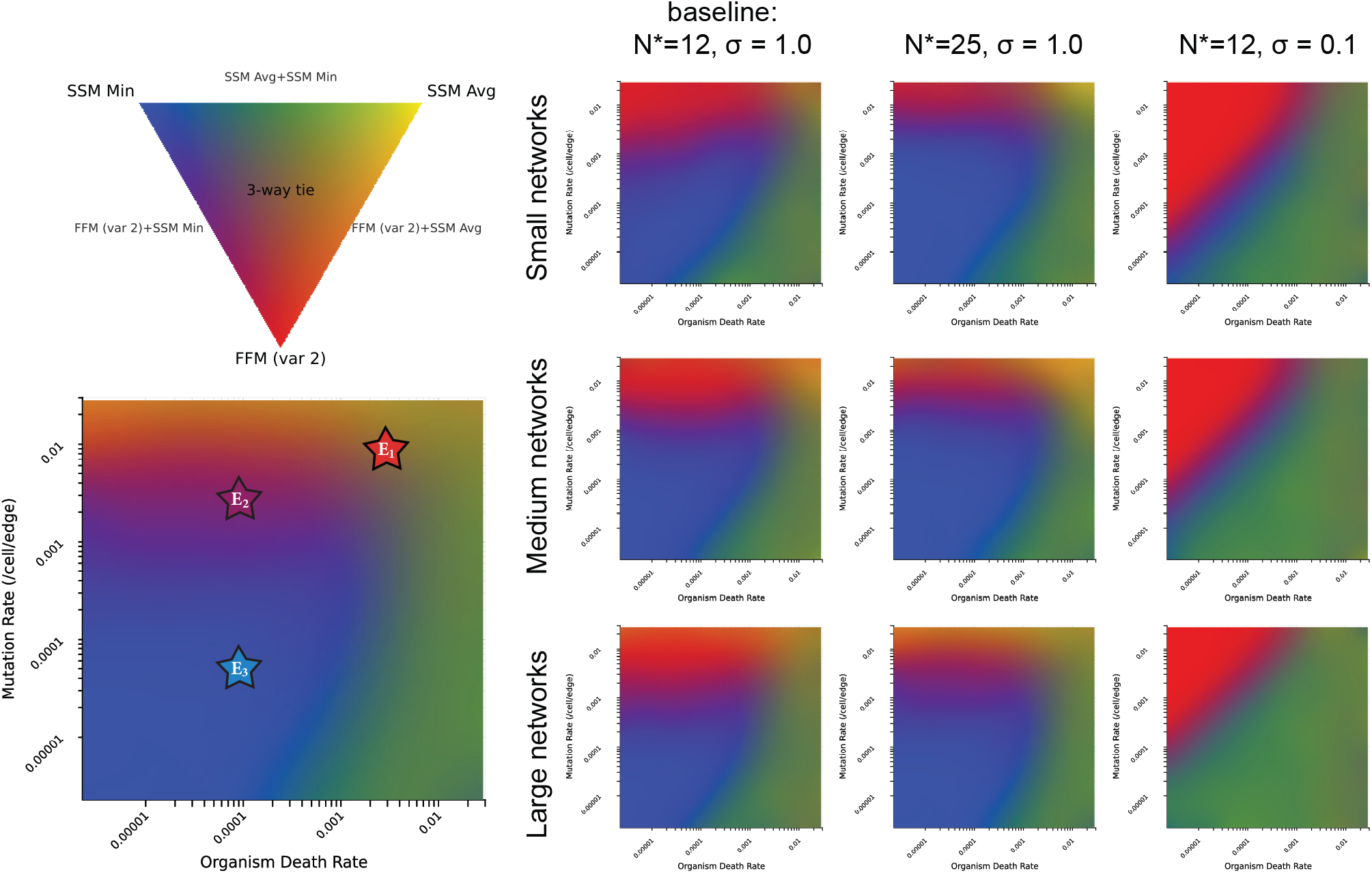
Each heatmap is a log-scaled grid of organism death rates and mutation rates and is colored indicating probability that each of the following three network architectures has the highest fitness for any size up to the maximum network size (80 mutatable parameters) according to an RYB color palette: feedforward motif averaging redundancy variant 2 (red), simple step motif averaging redundancy (yellow), and min redundancy (blue). Feedforward motif averaging redundancy variant 1 is not included in heatmaps due to low relative fitness everywhere. To calculate network architecture superiority in heatmaps, 20 replicates of fitness were collected in simulations of parsimonious creatures for each position in a 127×127 grid of mutation rates and organism lifespans and across all network sizes independently for all network types, where 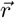 for each redundant network architecture and size was optimized for hedging utilizing a spatial evolutionary algorithm across the grid of environments. Bootstrapped t-tests were performed at each position in the grid of environments to calculate the probability that a particular network architecture has higher fitness than all other network types, then these three probabilities were colored at each position according to an RYB color gamut. The leftmost column of heatmaps are for the baseline case where *σ* = 1.0 and *N*^*\**^ = 12. Rows of heatmaps cut off maximum network size at small (8 parameters), medium (30 parameters) and large (80 parameter) networks, while the 2nd and 3rd columns indicate either an change from the baseline to *N*^*\**^ = 25 or *σ* = 0.1 respectively. A legend and an enlarged heatmap is shown indicating E1-E3 positions on this grid for comparison with other figures.

**Figure S4:**
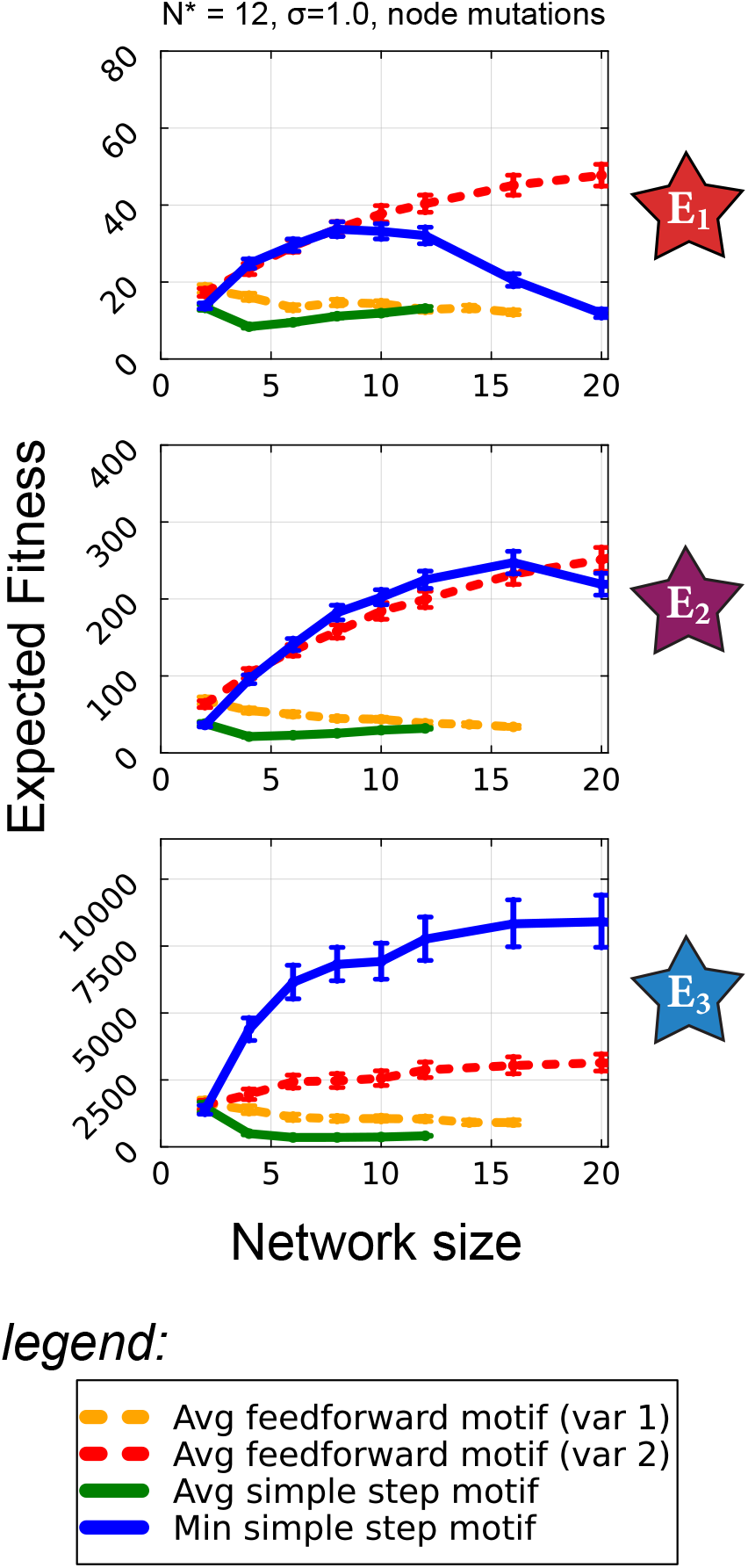
In each plot, expected fitness vs number of mutating parameters is plotted for feedforward averaging redundancy networks, ssm min redundancy and ssm averaging redundancy, in the baseline case where *N*^*\**^ = 12, *σ* = 1.0, but where mutations impact the rate of transcription factor production itself (node mutation) rather than interaction strengths between transcription factors (edge mutation) as was performed for data in the main text. As such, network size is measured as the number of mutateable parameters, which will be the number of nodes rather than the number of edges. Each plot focuses on a specific combination of organism death rate and mutation rate indicated by the stars *E*_1_, *E*_2_, and *E*_3_ (see text for details). To account for hedging, homogeneous 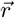 was optimized using Bayesian Optimization prior to estimating fitness of each network tested. Expected fitness was estimated by measuring fitness for many independent simulations of parsimonious creatures with the exact same optimized network, and 95 percent confidence intervals were estimated using bootstrapping.

**Figure S5:**
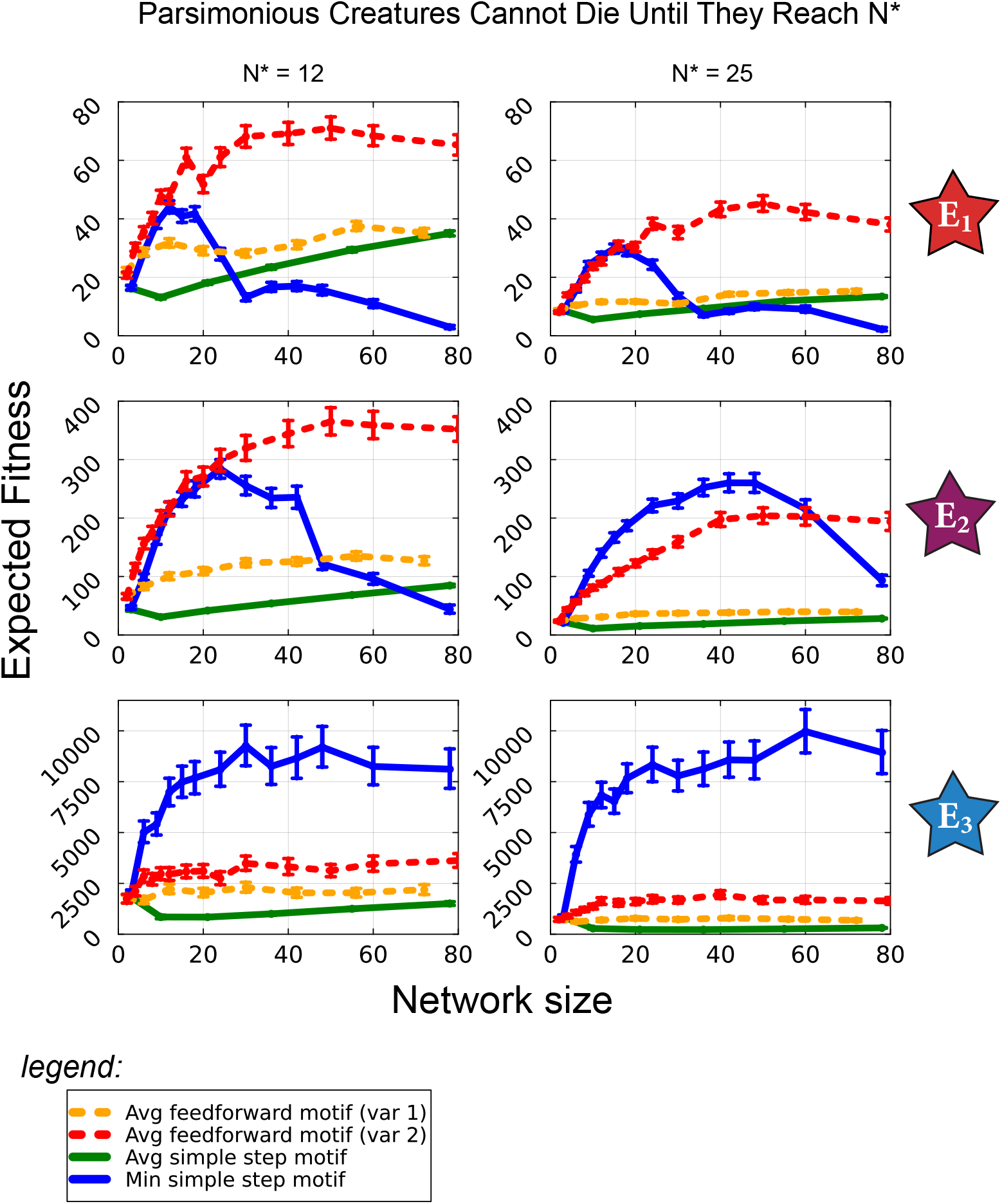
In each plot, expected fitness vs number of mutating parameters is plotted for feedforward averaging redundancy networks, ssm min redundancy and ssm averaging redundancy, but where each parsimonious creature simulated cannot undergo organism death from when they are born as a single cell until they have reached *N* = *N*^*\**^ for the first time, at which point organism death can begin to occur. Each plot focuses on a specific combination of organism death rate and mutation rate indicated by the stars *E*_1_, *E*_2_, and *E*_3_ (see text for details). The leftmost column of plots indicate E1-E3 where *N*^*\**^ = 12, *σ* = 1.0, and the rightmost column of plots where *N*^*\**^ has been increased to 25. To account for hedging, homogeneous 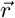 was optimized using Bayesian Optimization prior to estimating fitness of each network tested. Expected fitness was estimated by measuring fitness for many independent simulations of parsimonious creatures with the exact same optimized network, and 95 percent confidence intervals were estimated using bootstrapping.

**Figure S6:**
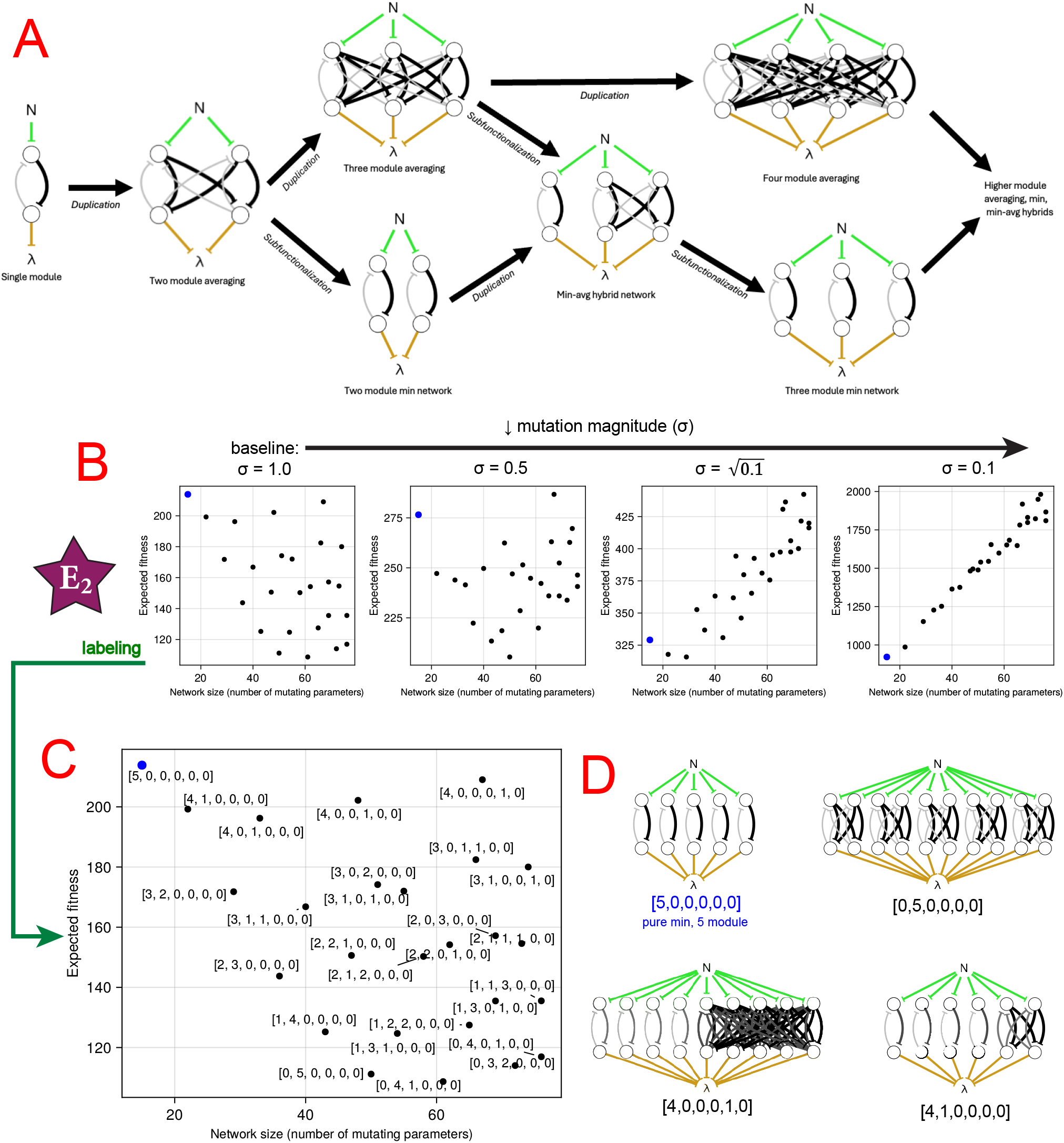
Possible pathways of formation of averaging redundancy, min redundancy, and min-avg hybrid networks, with comparison of expected fitness of a subset of min-avg hybrids with respect to network size. (A) A single simple step motif module may go through cycles of duplications and/or subfunctionalization to form redundant pathways to regulate the cell’s replication function while retaining a replication function nearly identical to that with a single motif. Immediately after a complete duplication of a module, homologous interactions will remain between modules, forming averaging redundancy. These homologous interactions must be severed to create fully independent modules, forming min redundancy, while min-avg hybrid networks form as intermediates. (B) Expected fitness vs network size for hybrid networks possessing 5 min() arguments. E2 is selected with decreasing mutation magnitude. To account for hedging, homogeneous 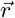 was optimized using Bayesian Optimization prior to estimating fitness of each network tested. Expected fitness was estimated by measuring fitness for many independent simulations of parsimonious creatures with the exact same optimized network, but 95 percent confidence intervals were not included as to avoid crowding plots. (C) Leftmost plot from B is enlarged and each hybrid network type is labeled as a vector encoding the degree of averaging in each module for which the minimum is taken. (D) Legend illustrating labeling of hybrid networks in C. Top left in blue shows an example encoding vector for a pure min redundancy network of 5 modules, examples of other min-avg hybrid networks are illustrated with their corresponding encoding vectors.

**Figure S7:**
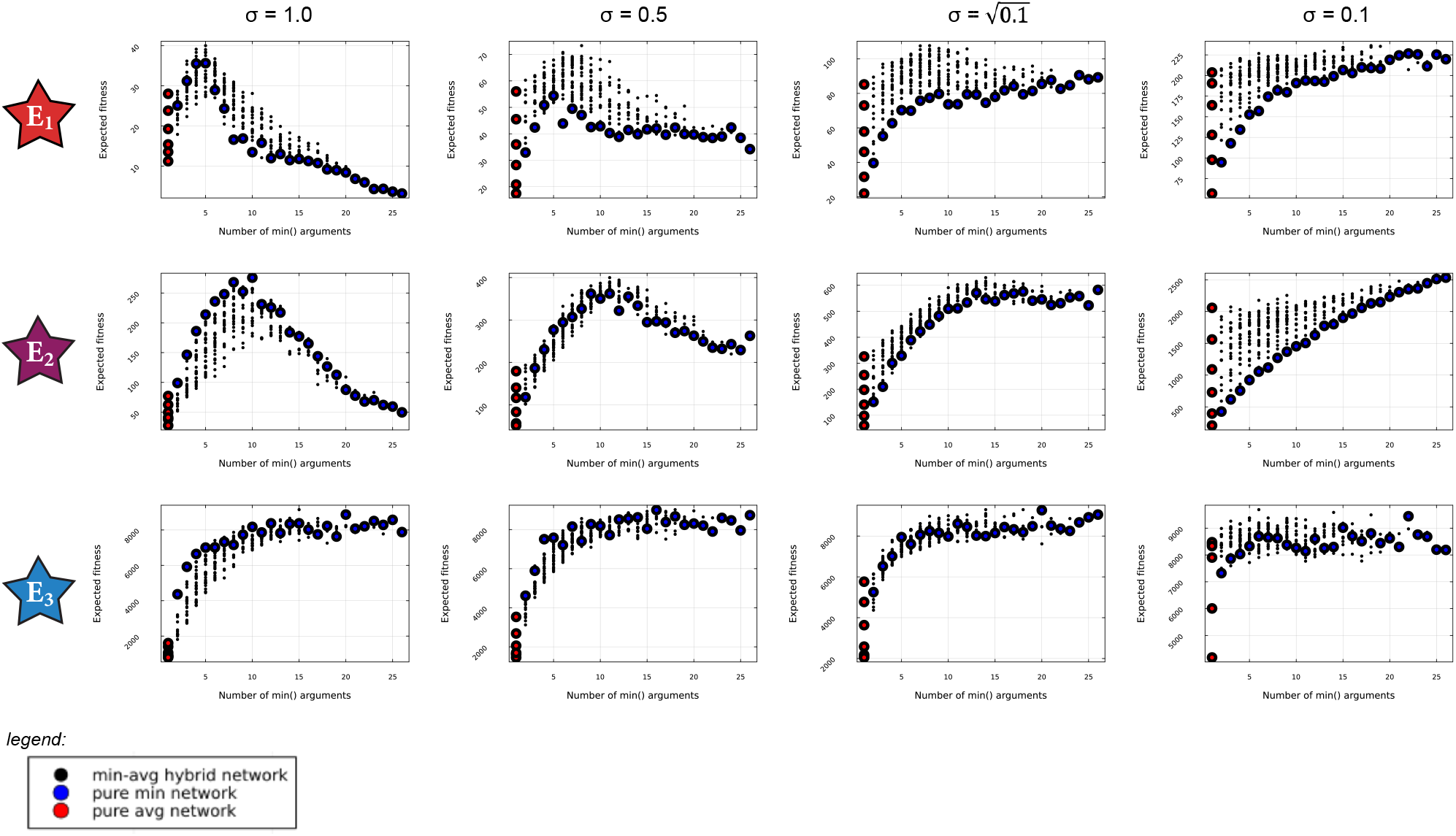
In each plot, expected fitness vs number of min() arguments is plotted for all 312 unique min-avg hybrid networks formed from the simple step motif up to 80 mutateable parameters. Where pure averaging redundancy is plotted in red, and pure min redundancy in blue. Each plot focuses on a specific combination of organism death rate and mutation rate indicated by the stars *E*_1_, *E*_2_, and *E*_3_ (see text for details). The leftmost column of plots indicate E1-E3 where *N*^*\**^ = 12, *σ* = 1.0, and columns of plots test decreasing *σ* from from left to right. To account for hedging, homogeneous 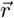 was optimized using Bayesian Optimization prior to estimating fitness of each network tested. Expected fitness was estimated by measuring fitness for many independent simulations of parsimonious creatures with the exact same optimized network, and 95 percent confidence intervals were not included as to avoid crowding plots.

